# Revisiting the auditory hemodynamic response function in the era of fast fMRI

**DOI:** 10.64898/2026.06.24.734180

**Authors:** Letitia Schneider, Isma Zulfiqar, Yaël Balbastre, Lori L. Holt, Martina F. Callaghan, Frederic Dick

## Abstract

Recent advances in fast fMRI now enable whole-brain imaging with a TR of ≤1 s, which has helped to rekindle interest in characterizing the blood oxygen level dependent hemodynamic response function (BOLD HRF). Recent studies in the visual system have found intra-areal differences in temporal response characteristics, as well as HRFs that were faster and narrower than predicted by standard models. The auditory system presents a unique challenge, in that neuronal populations must operate across timescales of microseconds to minutes, and the surface of auditory cortex in particular is intricately and heavily vascularized. Here, we used fast fMRI to characterise voxelwise auditory HRFs evoked by short naturalistic sounds, assessing HRF reproducibility and variability across sessions, participants, and independent datasets. Across two studies at 3 T, participants passively listened to short environmental sounds while fMRI and quantitative MRI data were acquired with a 1s TR. Voxelwise HRFs were estimated via novel cross-session alignment routine, and exclusion of large vascular contributions. We identified a diverse set of hemodynamically plausible response shapes, which were not consistently captured by standard HRF approaches. These responses were reproducible within participants across sessions and robust across two independent acquisitions. Using data-driven gamma models, we achieved stable estimates with relatively few runs, particularly in auditory temporal regions. Within auditory cortex, we observed reproducible spatial gradients in response timing and shape, with faster and higher magnitude responses in medial regions, and slower and lower magnitude responses laterally. Together, these findings demonstrate that auditory HRFs are diverse, reliable, and regionally specific, and highlight the value of fast fMRI and data-driven modelling for advancing interpretation of fMRI data.

## 1 Introduction

Hemodynamic response functions (HRFs) evoked by even simple stimuli can show markedly different shapes across brain regions, with some work suggesting that different components of the HRF may reflect distinct underlying neural processes, such as inhibition, a particularly important mechanism in auditory function (Chen et al., 2023; Szycik et al., 2012). Despite this variability, most fMRI analyses still use a standardized approximation of the response, adopting a canonical HRF as a practical default framework for modelling BOLD dynamics. This HRF model is typically based on a gamma function that peaks 4-6 s after stimulus presentation and rests on the assumptions of a linear system (Boynton et al., 1996). Critically, reliance on canonical HRF models can produce false negative results potentially obscuring our understanding of underlying neural responses (Chen et al., 2023; Handwerker et al., 2004; Prince et al., 2022).

Recent advances in receive coils, parallel imaging, and pulse sequence design offer an opportunity for improved characterisation of the HRF. ‘Fast fMRI’, allows imaging of the brain with repetition times (TRs) of ≤ 1000 ms, including for whole-brain imaging at 3T (Bollmann & Barth, 2021; Polimeni & Lewis, 2021). Leveraging these developments, recent work has shed light on intra-areal HRF differences (Gomez et al., 2024). Using a flickering stimulus with a changing luminance contrast at different frequencies (0.05-20Hz), Gomez et al. (2024) discerned a spatial pattern of peak times within primary visual cortex (V1), with faster responses concentrated in anterior V1 and slower responses in posterior V1. These differences explained more variance in temporal characteristics of the HRF than factors like vasculature or cortical curvature (Gomez et al., 2024). This aligns with previous reports of magnitude and timing differences in the BOLD response within V1 (Bailes et al., 2023; Kurzawski et al., 2022). Taken together, studies capitalising on faster sampling rates and higher spatial resolution reveal fine-grained differences in voxelwise HRF timings even within regions, suggesting that canonical HRF models may not accurately reflect underlying response details accessible with current imaging standards (Polimeni & Lewis, 2021).

While technological advances have expanded our understanding of the HRF and challenged canonical models, much of the recent work – like the early studies motivating the canonical HRF model – still relies on visual stimuli. Previous evidence suggests that responses within the auditory cortex appear to be consistent across sessions and within its subregions (Baumann et al., 2010; Gutschalk et al., 2010; Taylor et al., 2022). Taylor et al. (2018) set out to characterise the HRF to an audiovisual stimulus across cortical grey matter by decomposing average HRFs into principal components. Their results showed that most of the variance in the observed voxel time-series was explained by a stereotypical HRF, marked by a relatively stable initial peak, width and undershoot across participants (Taylor et al., 2018). However, work combining neuronal and biophysical HRF models has demonstrated voxelwise differences in HRF characteristics within auditory regions (Zulfiqar et al., 2021). Specifically, the caudal belt region was best explained by neuronal models with faster temporal dynamics, whereas the rostral belt region by those with slower temporal dynamics (Zulfiqar et al., 2021). Thus, variability in the auditory HRF may exist but might have been missed previously due to averaging of heterogeneous voxel responses (Taylor et al., 2018).

The present study aims to directly address the above-mentioned discrepancies by employing fast fMRI to characterise the voxelwise HRF to short, naturalistic auditory stimuli. By focusing on voxelwise responses and controlling for macrovascular contributions, we expect to unveil differences in HRF characteristics that may have been missed before and might hold important details about underlying cognitive processes. To avoid constraints imposed by the use of canonical HRF models, we propose a novel data-driven method to estimate HRF shapes. To probe the previously shown stability of the auditory HRF, we acquired data from the same participants across different time points. We also acquired additional independent data from the same participants with a modified design and image acquisition protocol. This allowed us to test how robustly the HRF can be identified across independent acquisitions with varying spatial resolutions. The data-driven HRF models in tandem with the high temporal resolution used in this study stand to reveal whether fine-grained differences in temporal response characteristics exist within auditory cortex. We further examined whether temporal differences relate to underlying degrees of myelination, given prior evidence linking myelin markers to cortical vasculature (Edwards et al., 2023).

## 2 Methods

### 2.1 Experimental design

The experiment was designed to identify and characterize the full auditory hemodynamic response function (HRF) using brief auditory stimuli. Participants were presented with 500 ms auditory stimuli separated by long interstimulus intervals (ISI >18s) to allow the BOLD signal to return to baseline between trials.

#### 2.1.1 Study overview

Each fMRI session lasted approximately 40 minutes, including a short structural scan, followed by six functional runs of ∼5 minutes during which participants passively listened to brief environmental sounds (Fig. 1). We acquired two independent fMRI datasets to compare HRF characteristics across acquisition conditions and to ensure observed effects were not protocol-specific. Variations in echo time (TE), bandwidth, and spatial resolution are known to alter BOLD contrast sensitivity, noise levels, and partial volume effects, thereby affecting the estimated amplitude, timing, and shape of the HRF (Buxton et al., 2004; Triantafyllou et al., 2005a). The first dataset (‘Original dataset’) consists of two fMRI sessions, and the second dataset (‘Replication dataset) consists of four fMRI sessions; each session contains six runs of the same experimental paradigm. The subset of participants who were common to both datasets were used to evaluate the stability of HRF estimates across protocols.

**Fig. 1.**
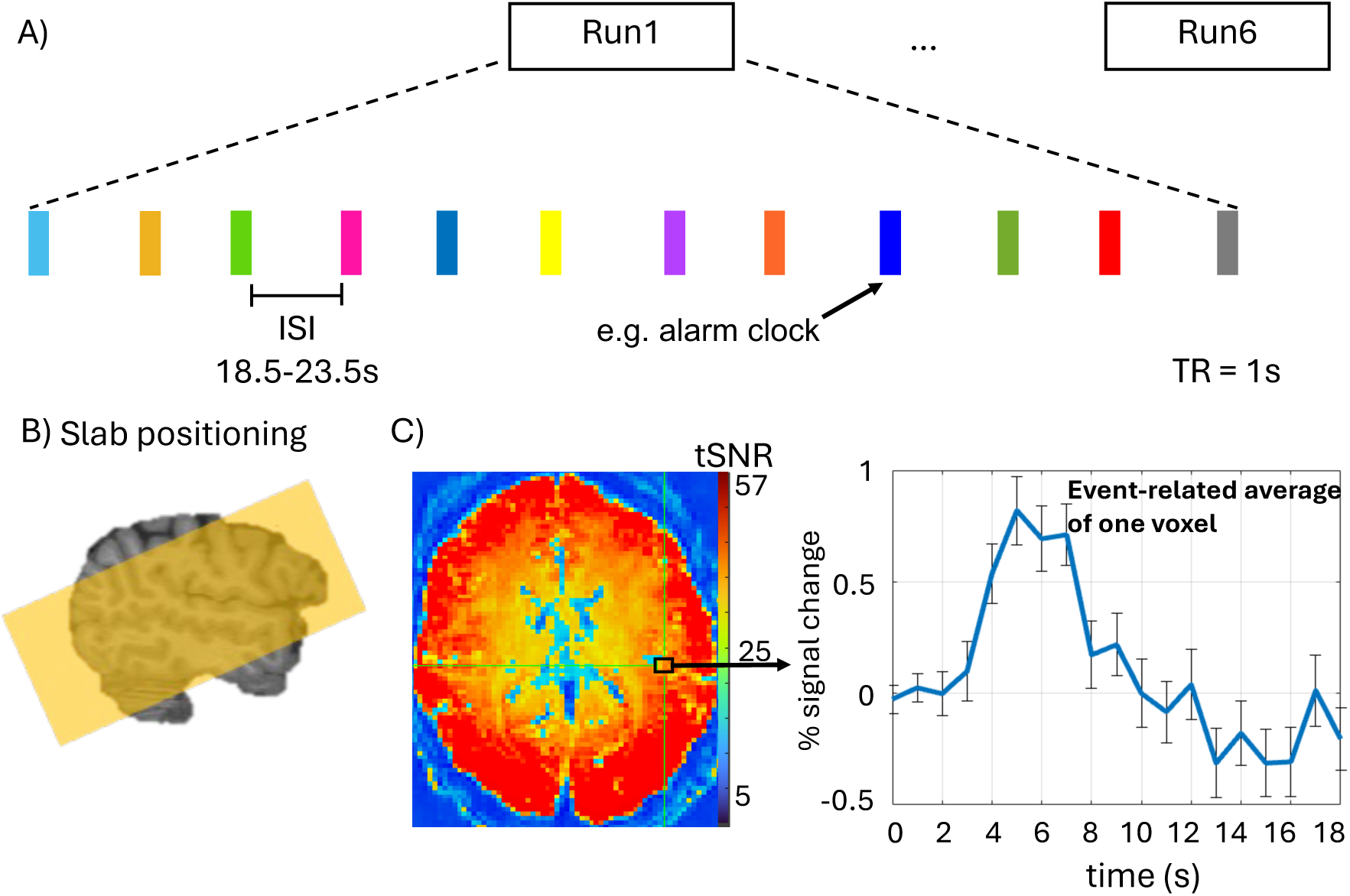
Experimental design and analysis. **(A)** Participants completed six fMRI runs per session passively listening to 500 ms auditory stimuli presented with long inter-stimulus intervals to allow for a characterisation of the full hemodynamic response. (**B)** The fMRI slab was centred on the transverse temporal gyrus bilaterally. (**C)** High tSNR voxels were identified from the tSNR map. Each of these voxel’s timeseries were averaged across sounds according to the stimulus onset times to create event-related averages.

#### 2.1.2 Participants

##### 2.1.2.1 Original dataset

Five healthy adults (three males; age 36 ± 10 years) with normal hearing and no previous history of neurological disorders participated in the study. The study was approved by the research ethics committee of University College London and participants provided written informed consent prior to the experiment. Each participant completed two fMRI sessions of six runs each, spaced no more than two weeks apart.

##### 2.1.2.2 Replication dataset

The Replication dataset included four participants from the Original dataset along with two additional adult healthy participants (two males; average age 35 ± 12 years), making a total of six participants. All participants provided written informed consent prior to scanning. Each participant completed four fMRI sessions of six runs each, with all sessions acquired within a maximum of 9 weeks.

#### 2.1.3 Stimuli and Design

##### 2.1.3.1 Original dataset

Each run consisted of twelve short environmental sounds, each lasting 500 ms (see example stimulus in Fig. 1A). Stimuli from Zhao et al. (2019). Twenty seconds of silence were added at the beginning and end of each run as a baseline measure. The interstimulus intervals varied between 18.5 and 23.5 s. These long intervals were chosen to ensure the response can return to baseline before the presentation of the next stimulus. Stimulus order was randomised across six runs, and each run lasted approximately four minutes 38 seconds.

##### 2.1.3.2 Replication dataset

The experimental design for the Replication dataset was identical to the Original dataset, except the ISI was fixed at 19.58 s. This ISI was selected to accommodate the requirements of a parallel experiment conducted within the same scanning session, while remaining well within the range shown in the Original dataset to be sufficient for reliable HRF estimation. Participants listened to the same environmental sounds for six runs of approximately four minutes each, with a total of four sessions per participant.

### 2.2 MRI acquisition and preprocessing

#### 2.2.1 Stimulus presentation

For both datasets, stimuli were presented using a Mac mini via PsychoPy (v. 2022.2.4). Participants heard the sounds through custom MRI-compatible earphones (‘OMEMS’, Josephs et al., 2025) connected to an external amplifier (DENON PMA-255UK). Prior to the first fMRI session, the sound volume was adjusted individually to ensure stimuli could be heard comfortably above scanner noise. Anatomical scans were acquired first to aid slab positioning for subsequent EPI acquisitions. Additionally, five EPI volumes with inverse phase encoding direction (P->A) were acquired after the first run for geometric distortion correction. Slab positioning angle was kept constant between session, and participants were placed in a novel head stabilisation device within the head coil, (MR-MinMo, patent pending, UK (GB20220005139|WO2023GB50932), US (US2025213412), Europe (EP4504061) and Japan (JP2025511707). The device minimises both intra-session motion and inter-session variance in head position (Mangal et al., 2026, Levchenko et al., 2026).

#### 2.2.2 Structural image acquisition

All MRI data were acquired on a 3T Siemens Prisma scanner. Structural scans for both datasets were T1-weighted anatomical images used for slab positioning and surface reconstruction. These were 3D magnetisation-prepared rapid acquisition gradient-echo (MPRAGE) sequence (GRAPPA acceleration factor 4) with a voxel size of 1 × 1 × 1 mm (repetition time (TR) / echo time (TE) / Inversion Time = 1530 / 2.98 / 900 ms, flip angle = 9 degrees, matrix size = 256 × 256 x 176). In a subset of participants (see above), multi-parameter mapping (MPM) (Weiskopf et al., 2013) were also acquired in a different session to estimate proxies of myelination. Proton density-weighted (PDw), T1-weighted (T1w), and magnetisation transfer (MTw) images were acquired using an <u>in-house</u> multi-echo 3D FLASH pulse sequence (voxel size: 0.8 x 0.8 x 0.8 mm, matrix = 320 x 280 x 224, TR = 25.0 ms, bandwidth 490 Hz/px,), slab rotation 30°, excitation flip angle: 6° (PDw/MTw) or 21° (T1w). Participants were instructed to fixate on a fixation cross during acquisition to minimise eye motion artefacts. Total acquisition time for the MPM protocol, including calibration data to map transmit and receive fields, was approximately 28 minutes.

#### 2.2.3 Functional image acquisition

After collecting the T1-weighted image, sagittal-oblique slices were positioned based on this image to target the transverse temporal gyrus for subsequent BOLD-weighted functional runs. Importantly, functional images were acquired with a relatively short TR of 1s at 3T to achieve higher temporal resolution when sampling the HRF. To achieve the study aim of characterizing the variability of the HRF across cortex, the field of view included 48 axial slices providing near whole-brain coverage across participants.

##### 2.2.3.1 Original dataset

Functional images were acquired using gradient echo-planar imaging (EPI) sequences written and supplied by the Centre for Magnetic Resonance Research (https://downloads.cmrr.umn.edu/software/package/multiband/) (Moeller et al., 2010), with 4-fold multiband acceleration (Feinberg et al., 2010), 2mm isotropic resolution, TR = 1000 ms, TE = 35.2ms, flip angle = 60 degrees, field of view = 212mm. Bandwidth was 2620 Hz/px and slices were acquired in interleaved order.

##### 2.2.3.2 Replication dataset

Functional images were acquired using the same multiband EPI framework (Moeller et al., 2010; Feinberg et al., 2010), also with 4-fold multiband acceleration. To evaluate whether HRF estimates generalise across acquisition settings, images were collected at 2.5 mm isotropic resolution with a 0.5 mm slice gap, TR = 1000 ms, TE = 28 ms, flip angle = 60°, FOV = 210 mm, 6/8 partial Fourier acquisition, and bandwidth = 1985 Hz/px, (see summary in Supplementary Table1). Slices were acquired in ascending order.

#### 2.2.4 Functional data preprocessing

Imaging data from both datasets were processed in the same way. Functional images were converted from DICOM to NIFTI format using dcm2niix (Li et al., 2016) and preprocessed using afni_proc.py in the AFNI software (v. 23.0.02, Joo et al., 2010). The first eight images were discarded to allow for longitudinal magnetisation to reach equilibrium. EPI data were unwarped using the single band image from the first run (= blip-forward dataset) as well as from the subsequent single-band image from the phase-reverse-encoded sequence (= blip-reversed dataset). Single band images were used for unwarping because of their higher tissue contrast. Motion correction was performed by aligning all functional runs within a session to an image that detrending determined to be least different from all other images (i.e., minimum outlier volume) using volreg and regress blocks in afni_proc.py. The motion reference EPI volume was registered to each participant’s reconstructed cortical surface (v 7.3.2, Dale & Sereno, 1993) (v 7.3.2, Dale & Sereno, 1993) after aligning it to average-space (see 2.2.6).

#### 2.2.5 Structural data preprocessing

Data were processed using the hMRI toolbox in SPM (Tabelow et al., 2019). This produced a map of longitudinal relaxation rate (R_1_= 1/T_1_). A synthetic MPRAGE-like image was created based on scaled R_1_ and proton-density maps using mri_synthesize in Freesurfer. This synthetic image was then processed using FreeSurfer’s recon-all pipeline, which performs cortical surface reconstruction in FreeSurfer’s native 1 mm anatomical space, including skull stripping using the proton-density volume (Dale & Sereno, 1993; Dick et al., 2017). Reconstructed surfaces were visually assessed for quality and used to derive anatomical regions of interest (ROIs; see 2.3.7. for more details). Quantitative R_1_ maps were projected into the surface using the Freesurfer program *mri_vol2surf* and sampled at 5 points between 30 to 70% of the cortical depth fraction. Using *mris_anatomical_stats* ROI-wise R_1_ estimates were extracted from each participant, and hemisphere and outputs were saved as structured text tables for downstream analysis.

#### 2.2.6 Cross-session registration

In addition to placing participants in the MR-MinMo head stabilisation device and keeping slab positioning constant, novel cross-registration tools were used to facilitate more precise cross-session comparisons. One goal of the current study was to characterize the degree of HRF similarity and stability at as high a granularity as possible, ideally at the voxelwise level, and with as little interpolation and smoothing as possible across different tissue types (including blood vessels). This is challenging when EPI images have been acquired across two or more separate sessions (or when the head has moved significantly), because of the session-specific nonlinear and local geometric image deformations caused by the imperfectly shimmed B0 field. To address this problem, we used a symmetric diffeomorphic alignment procedure inspired by Casamitjana et al., (2025), which aims to find an optimal average space via a series of pairwise registrations. Briefly, given a set of images acquired during different sessions, this method consists of linearly and nonlinearly co-registering all possible pairs of images. It then leverages the Lie group representation of both the linear and nonlinear transformations to infer an optimal mean space across all sessions, along with a set of transformations mapping from this mean space to each session. Time-series from all sessions can then be resliced into this mean space, which avoids biasing the analysis towards either session due to non-symmetric interpolation artefacts. While similar to Casamitjana et al., (2025) in concept, our implementation differs in its details and is described in Appendix 1. In practice, the algorithm was applied to the EPI native-space minimum outlier volumes from each session, and zeropadded by 6 voxels at superior and inferior edges. This zeropadding step is commonly recommended in AFNI workflows as it preserves spatial correspondence across sessions.

#### 2.2.7 Creation of tSNR mask and surface projection

For each motion- and distortion-corrected functional run, data were first detrended run-wise by removing the mean and slope of each voxel time-series. Temporal signal-to-noise (tSNR) maps were then computed by dividing the mean of each time-series by its standard deviation. The average tSNR map for each session was zeropadded first and then resliced into the average-space of all sessions using the linear and nonlinear transformation outputs from the co-registration output (see above). To estimate and exclude voxels that likely lay in large vessels (Moerel et al., 2018; Zhang et al., 2009), binary masks were generated from the average-space-aligned tSNR maps based on a visually determined threshold (tSNR=35 for all participants), with reference to known patterns of vasculature (Duvernoy et al., 1981). To further restrict the voxels in the mask to the brain, *3dAutomask* in AFNI was applied to the average-space-aligned motion reference EPI volume, and the conjunction of this mask with the tSNR mask was computed.

Following the same procedure used for the average tSNR map, motion- and distortion-corrected images for each session were zeropadded and resliced into average-space. Using FreeSurfer’s *mris_volsmooth* function, average-space EPI data from each frame were sampled at 3 points between 20-80% of the cortical depth fraction onto the cortical surface, smoothed along the surface using a 2 mm 2D FWHM kernel, and the average values across the cortical depth fractions were then projected back into the average-space EPI volume. Lastly, timeseries for each voxel within the tSNR mask were used to generate event-related average responses across all trials in MATLAB (Version 9.13, R2022b).

### 2.3 HRF identification

#### 2.3.1 Event-related response extraction

Voxelwise event-related responses were computed for the 72 trials (12 trials per run, 6 runs in total) for each session. The responses were estimated using a time window ranging from 0 (onset) to 18s post onset and were normalised to percent signal change relative to baseline signal (the average of the three TRs comprising time points before, at and immediately after stimulus onset). Normalisation was performed by subtracting the baseline signal from each time point and dividing the result by the baseline. To further remove noisy voxels from the data, additional quality checks were performed across all sessions. Firstly, low-intensity voxels (< 1500 in arbitrary units determined by the scanner) were excluded, because these were mostly on the border of the brainmask. Voxels with extreme event-related responses (based on 0.01st and 99.99th percentile cut-off across all participants) at any time point over any trial were removed. Event-related responses were computed from the voxels that passed this threshold and were common to all sessions. At each timepoint of these stimulus-locked time windows, the standard deviation was calculated across trials, yielding a trial-to-trial variability estimate for that voxel and timepoint. Voxels with a standard deviation greater than 3 at any time point (i.e., 95th percentile across all participants) were removed. Next, the mean correlation between the timecourses associated with all 72 sounds was computed per voxel, and a distribution across participants was generated. Voxels with mean correlations below the 85th percentile (a Pearson correlation of 0.02) were removed. For each of the remaining voxels, the time courses were averaged across runs within each session to generate event-related averages (ERAs, see Figure 1C).

#### 2.3.2 Hierarchical clustering of ERAs

Hierarchical clustering was employed as an exploratory, data-driven approach to identifying distinct HRF response shapes without imposing prior assumptions about their structure. Clustering was performed on the cleaned ERAs from the Original dataset Session_1 data (data from all participants was concatenated) using JMP, Version 17.0 (SAS Institute Inc., Cary, NC, 1989–2023). Within the Original dataset, Session_1 data was used as “training” data so Session_2 data could be used to evaluate the model accuracy (see 2.3.5). Voxelwise data were submitted to clustering using the *robust standardisation* option in JMP, which minimises the influence of outliers and reduces bias towards any single feature (e.g. amplitude). The total number of clusters was set to 40 to ensure that a wide variety of HRF shapes could be captured. Of the 40 identified cluster means, five were excluded due to high variability in the cluster mean or failure to return to baseline. A total of 40 clusters was selected because lower cluster numbers failed to capture the full diversity of hemodynamically plausible response shapes observed in the data, risking an oversimplified characterisation of HRF variability. For clarity of visualisation and to facilitate systematic visual comparison, the remaining 35 clusters were organised into higher-level cluster groups based on visual inspection of similarity in their shape profiles (Fig. 3).

#### 2.3.3 HRF curve fitting

The cluster means were modelled using one, two or three gamma functions, with the HRF expressed as a linear combination of these gamma functions (Worsley et al., 2002):

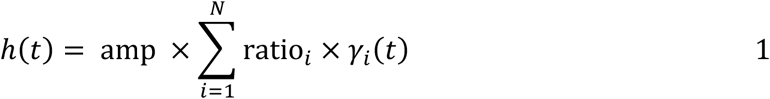

where *amp* denotes a global scaling parameter and ratio_%_ are relative weights, with *ratio*_1_ = 1 for identifiability. The maximum number of gamma components (*N*) was fixed at 3 (based on the identified HRF shapes).

Each gamma function *γ_i_* is the unnormalised density of a Gamma distribution, defined as (Proulx et al., 2014):

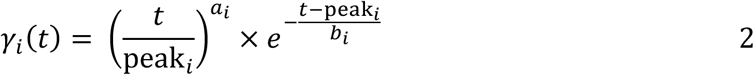

with parameters:

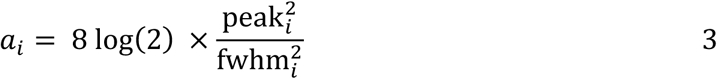

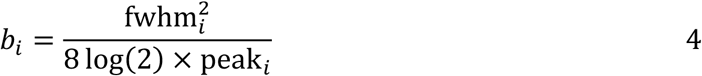

where peak_i_and fwhm_i_ correspond to the time-to-peak and full-width at half-maximum (FWHM), respectively. Each basis function is normalised to attain unit amplitude at its peak whereas amp scales the whole HRF (Proulx et al., 2014). Two- and three-gamma models (Eqs. 6 and 7) included additional parameters for peak amplitude ratios. The model dynamics were defined as follows:
1- Gamma model:

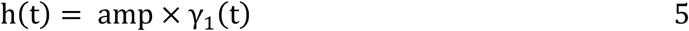

2- Gamma model:

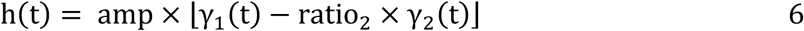

3- Gamma model:

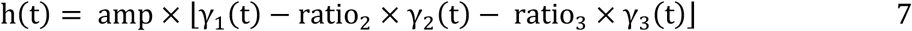

where ratio_2_ and ratio_3_ control the relative contributions of secondary and tertiary components respectively. Three-gamma models fell into two categories. The first consisted of one positive peak, followed by an undershoot and a later positive peak (with early and late variants). The second began with a fast negative peak (i.e. initial dip response), followed by a positive peak and a third peak of variable polarity (see Fig. 2).

**Fig. 2.**
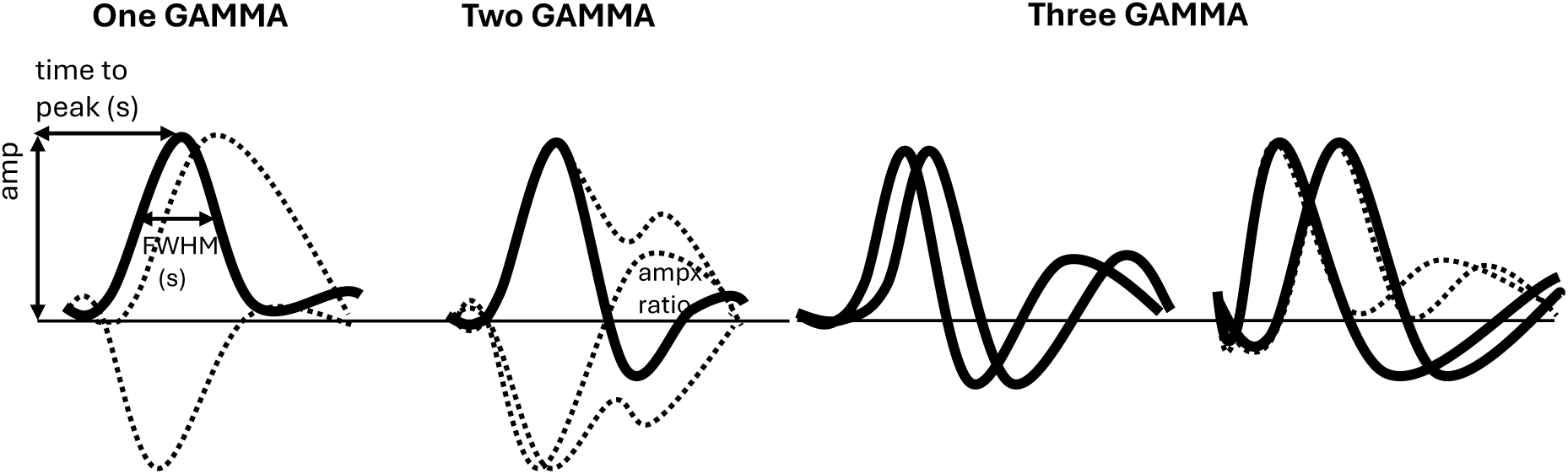
The observed shapes were summarised into six flexible, hemodynamically plausible models. Illustration shows the six data-driven models: one model with a single gamma function, one model with the sum of two gamma functions, and four models comprising three gamma functions each. Dotted lines represent different variants of the models. The shape of the gamma components is determined by the formula shown where peak controls peak latency and FWHM controls width.

#### 2.3.4 Voxelwise model fitting and validation

The 35 clusters were visually inspected and categorised based on their peak shapes and widths. These informed the lower and upper bounds of the model, as well as the starting points. Two checks were applied to each model fit to avoid overfitting to noise. First, the distance between consecutive peaks was required to be at least 2s. Second, the amplitude of all peaks was required to exceed 0.05 % signal change, ensuring sufficient activation in a voxel. An exception was made when modelling the initial dip (in two- and three-gamma models), whose amplitude threshold was set to 0.01% of signal change. Model fitting was performed using the nonlinear least-squares optimisation implemented in MATLAB’s *fit* function. Adjusted R^2^ (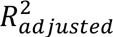) was used to quantify model fit, accounting for the varying number of fitted parameters across models (https://www.mathworks.com/help/stats/coefficient-of-determination-r-squared.html):

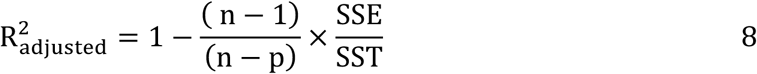

where SSE is the sum of squared error, SST is the sum of squared total, n is the number of observations, and p is the number of regression coefficients.

#### 2.3.5 HRF model comparison using cluster means from independent data

To assess how well existing HRF models capture the responses derived from each cluster, we systematically compared the goodness of fit (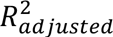) for each cluster mean time-series across different basis function sets. Since only Session_1 data from the Original Dataset were used to generate the models, Session_2 data were used here as independent data to evaluate model accuracy. Namely, cluster means obtained from hierarchical clustering on the cleaned ERA of Session_2 were fit with multiple models for comparison (see 2.3.2 for more details on clustering).

As an example of a constrained model, we included the canonical two gamma model from SPM (Friston et al., 1998). The canonical HRF from SPM was created in MATLAB using the default parameters of *spm_hrf* with a 1s TR. A stick function representing stimulus onset was convolved with the HRF and the resulting predictor was fit to the observed data using least-squares to estimate a scale (amplitude) and baseline offset. 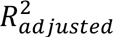 was then computed for each cluster mean. To increase flexibility, we also tested an extended version of the canonical model that included dispersion and temporal derivatives (Friston et al., 1998). This resulted in three basis functions which were combined before fitting; the combined predictor was again fit using least-squares scale + baseline estimation.

Next, for an even more flexible model compared to the canonical SPM HRF model with derivatives, we used the GLMsingle basis set, which contains a library of empirically derived basis functions (Prince et al., 2022). Each of the 20 HRFs from the *glm_single* library was convolved with the same stick function as above. For each resulting predictor, a scale and baseline term were fitted to the cluster mean, and the basis function yielding the highest 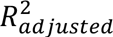 was selected as the best GLMsingle fit for that cluster mean. Finally, each cluster mean was also fit with each of the six custom gamma-based models developed from Session_1 data. For fitting, the same checks as outlined in 2.3.4 were performed to avoid fitting to noise, and overfitting. If these checks were unsuccessful for all six models, the cluster mean was excluded from further analysis. Otherwise, the model with the highest 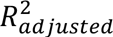 was selected as the best-fitting model. For the selected model, the HRF curve was reconstructed using its estimated parameters and convolved with the same stick function as fit to the data with least-squares scale + baseline estimation. Then the 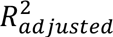 was recomputed for this convolved predicted timecourse. The time-series from the 40 clusters identified from Session_2 data were fit to both the custom models and standard HRF models of varying flexibility. Four clusters were omitted from further analyses because quality checks indicated that they had not resulted in a reliable fit to any of the six custom models. For each cluster, 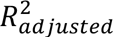 values from the competing models were compared using paired Wilcoxon signed-rank tests to account for within-cluster pairing. P-values were Bonferroni-corrected for the number of planned comparisons (custom model vs canonical HRF, canonical HRF with derivatives, and GLMsingle).

#### 2.3.6 Dynamics across runs – Simulation and run-wise stability assessment

Next, we asked how consistent HRF estimates are over time, and how much data are required for the HRF to reach relative stability. To examine HRF estimation over time (i.e., runs), we simulated data for 1000 voxels by assigning each to one of the predefined HRF shape classes (see Fig. 2 for shapes). Within each class, model parameters (i.e. peak latency) were sampled from specific ranges. Zero-mean Gaussian noise was added to the data to mimic real fMRI signals. This noise was informed by participants’ data, by using the trial-to-trial standard deviation computed for each voxel at each TR. For each voxel, the simulated and actual observed responses were averaged for different run combinations (i.e., Run_1, Run_1+2, Run_1+2+3…), and then fitted to each of the six models. The best fit (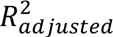) per voxel was recorded, averaged across all voxels, and plotted for each run combination, ranging from run 1 to run 6 (corresponding to Session_1) and up to run 12 (representing the average of all trials from both sessions) (Fig. 5). In the case of the Replication dataset run 24 corresponded to the average of all four sessions. To examine how responses evolve in Session_2 and whether adding an additional session improves stability of HRFs, model fits were also evaluated across the observed data of all runs only in Session_2. This analysis was performed separately for each participant. To identify, on a voxelwise level, which brain voxel model fits converge early (i.e., with fewer repetitions), we compared the 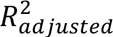 for each run to the maximum value obtained when all runs from Session_1 in the Original dataset were averaged. For each voxel, we tracked the 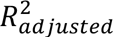 of the model that performed best with the full dataset (i.e. six runs) and determined the earliest run at which its 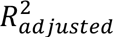 fell within ±20% of this final value. This allowed us to map the spatial distribution of response stability per voxel across the brain for each participant.

#### 2.3.7 ROI-based HRF characterisation

To test whether fast fMRI can reveal subtle differences in temporal properties of the HRF in auditory regions, we extracted model parameters corresponding to the first positive peak. Auditory regions of interest (ROIs) were generated based on the reconstructed cortical surface output from FreeSurfer *recon-all*. Labels from Sereno et al. (2022) were transformed from their original *fsaverage* space to each participant’s native space using Freesurfer’s *mri_surf2surf* tool. A total of 22 labels corresponding to auditory regions were extracted from the annotation using *mri_annotation2label*. The labels were then converted into volumetric space aligned to each participant’s EPI space using *mri_label2vol*, sampled from 20-80% of the cortical depth fraction and thresholded such that a voxel was included only if ≥50% of its volume was filled by the label. Finally, AFNI’s *3dmaskdump* was used to extract coordinate information from the volumetric ROIs. Three ROIs were excluded because they did not map accurately onto the cortical surface in any participant, leaving a final total of 19 ROIs per participant per hemisphere.

Several quality checks were performed before extracting temporal HRF features from the ROIs. First, if voxels were shared between ROIs, they were removed from the larger ROI. Furthermore, only ROIs containing more than five voxels were included in further analysis. ERAs were averaged across voxels, repetitions and sessions to produce one ERA per ROI. Each ROI ERA was fitted using the same methods and quality checks as for the voxelwise and cluster means fitting described above (see 2..3.4). ROIs with poor fits (<0.2 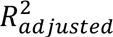) were excluded from further analysis to avoid misinterpretation of model estimates. Twenty-seven out of 412 ROIs, aggregated across both hemispheres, participants, and datasets, were excluded from analysis based on the criteria described above.

From the best-fitting model, key response characteristics were extracted for the first positive peak, namely peak latency, FWHM, and amplitude. These parameters were stored for each ROI for subsequent group-level analyses. The extracted information was used to gain insight into the distribution of these temporal HRF characteristics in auditory regions within and across participants.

In addition to assessing inter-individual differences in these temporal HRF characteristics, ‘delay and amplitude maps’ were generated for all participants across the two datasets. First, to evaluate general spatial patterns of peak timing and amplitude, the respective values obtained from the best model were averaged across hemispheres and normalized within each participant by subtracting the mean across that participant’s ROIs and dividing by the corresponding standard deviation. Normalisation was done to account for global inter-individual differences in peak delays and amplitude (see supplementary Fig. 1 for individual variability in HRF temporal characteristics). ROI-wise, within-participant z-scores were then averaged across participants for each dataset. Participant-averaged values were then projected on the *fsaverage* FreeSurfer surface for both datasets.

For the participants that were common across both datasets (n=4), we investigated the stability of ROI-wise peak latency, width, and amplitude across the two datasets using Pearson correlation. This was assessed in R (Version 2023.09.1+494; Posit team, 2025) and visualised in a scatterplot.

#### 2.3.8 Investigating the relationship between HRF shape and myelination

Given the known myeloarchitectonic profile of these auditory ROIs, we aimed to determine whether a relationship exists between temporal HRF characteristics and R_1_ – a cortical marker of myelination (Dick et al., 2017; Glasser et al., 2016; Lutti et al., 2014; Sereno et al., 2022). A linear mixed-effects model was fitted to assess whether local differences in R_1_, used as a proxy of myelination, were associated with differences in temporal response characteristics. This analysis was performed in R, accounting for hemisphere and participants-level variability. The model included fixed effects for hemisphere and R_1_, as well as a random intercept and slope for each participant:

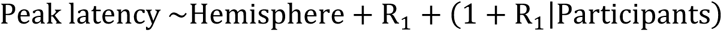

The dependent variable was either peak latency or amplitude, whereas the predictor was R_1_. Model estimates and significance were evaluated using summary statistics, with effective degrees of freedom approximated using the Satterwhaite method, as recommended for mixed models (Kenward & Roger, 1997). To visualize the relationship, scatterplots were generated using *ggplot2* in R.

## 3 Results

### 3.1 Hemodynamic responses to a short sound are characterised by different shapes

Clustering of Original Dataset Session_1 event-related averages revealed multiple HRF shapes associated with short sounds (see heatmap in Fig. 3). Five cluster means were excluded due to noise, yielding 35 cluster means that reflected hemodynamically plausible responses. These were observed across the cortex providing further support for previous evidence showing whole brain activation in response to even simple stimuli (Gonzalez-Castillo et al., 2012) (Fig. 4).

**Figure 3.**
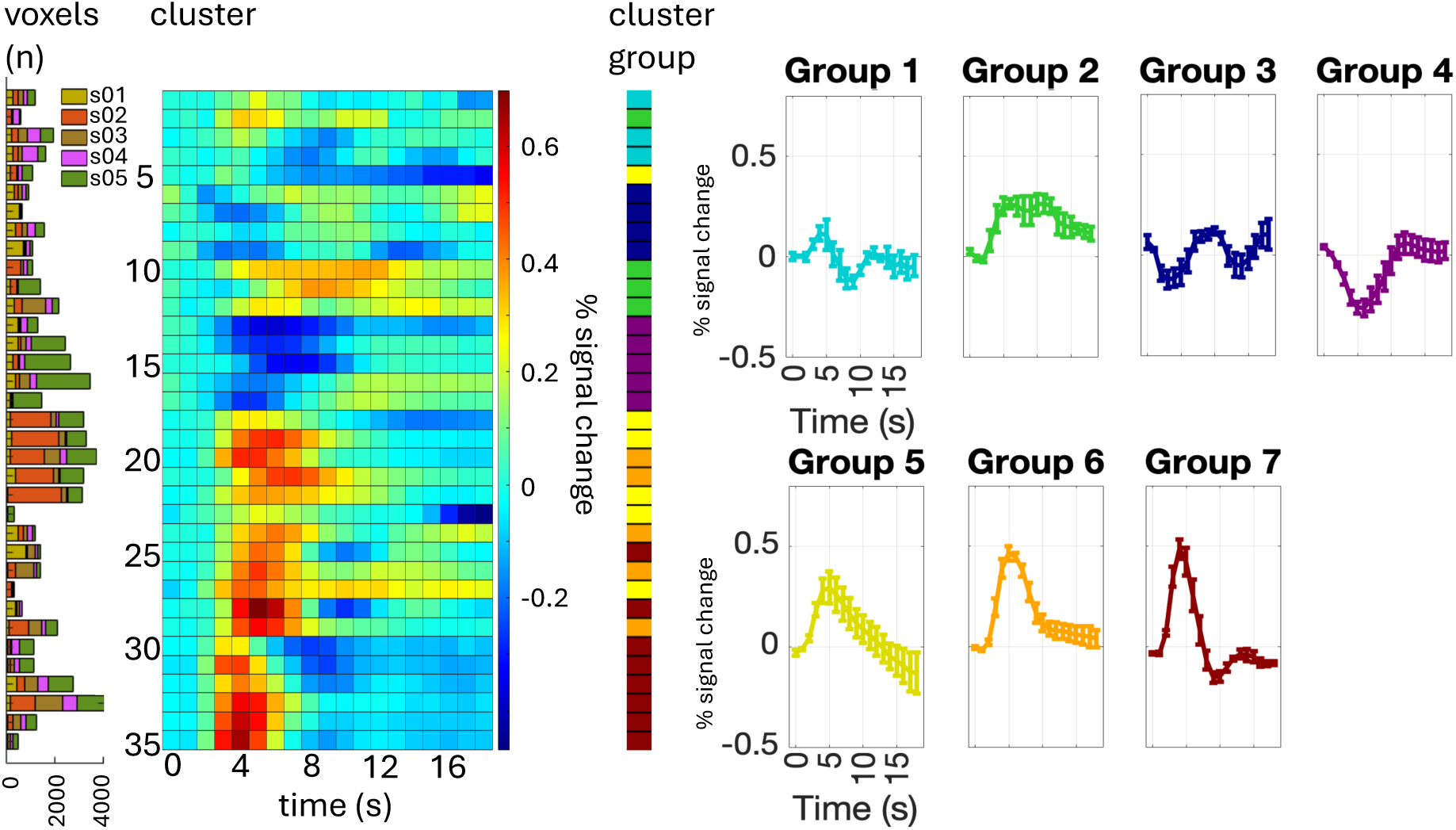
Listening to a short auditory stimulus evokes distinct voxelwise HRF shapes. Hierarchical clustering of Original Dataset Session_1 responses identified 35 clusters (matrix rows) each exhibiting hemodynamically plausible shapes. The colour bar indicates percent signal change. The number of voxels per participant per cluster are also shown (far left). These clusters were further summarised into seven distinct cluster groups which largely preserved the original clustering (far right). Error bars represent the standard error of the cluster means.

**Figure 4.**
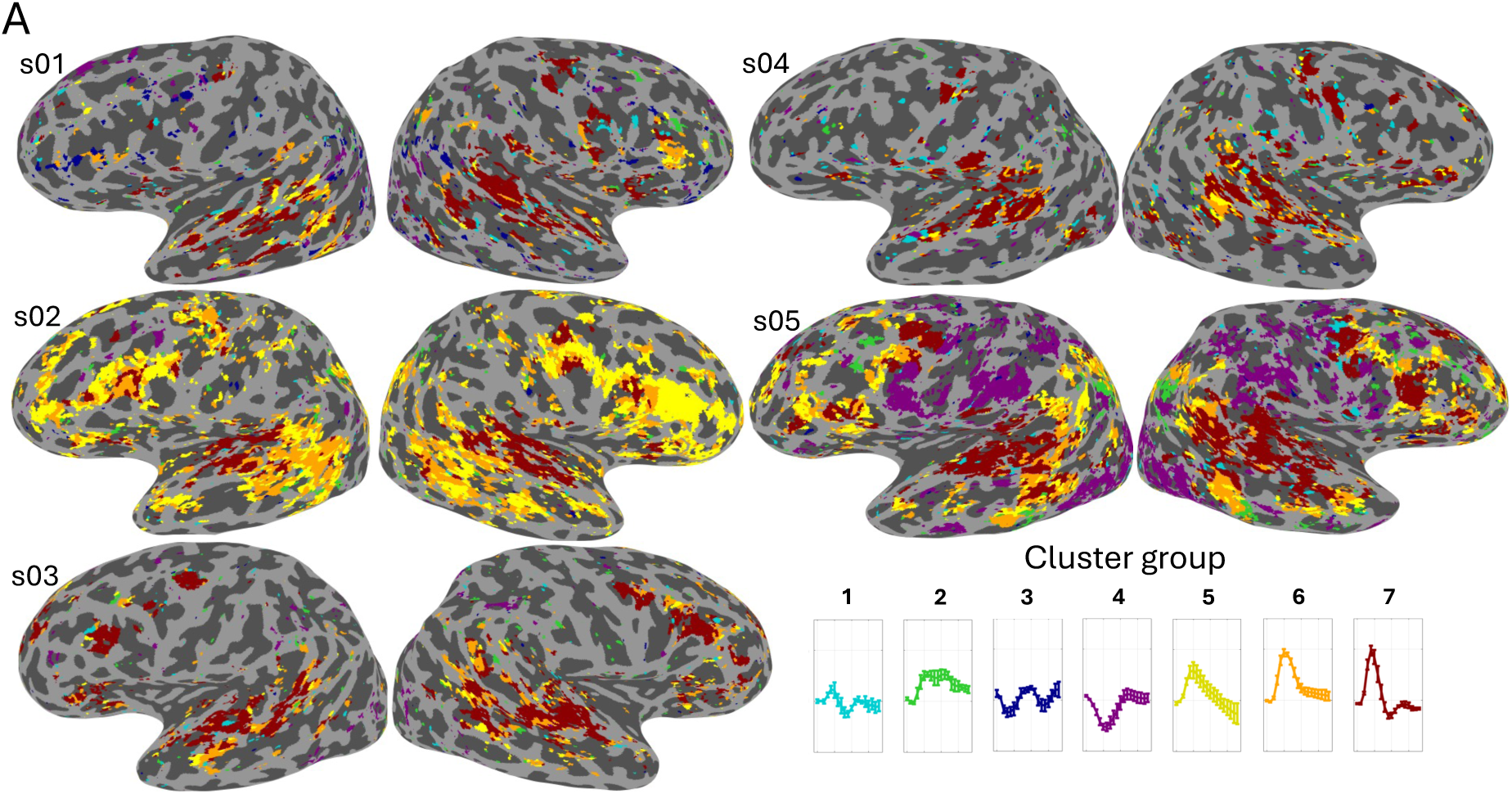
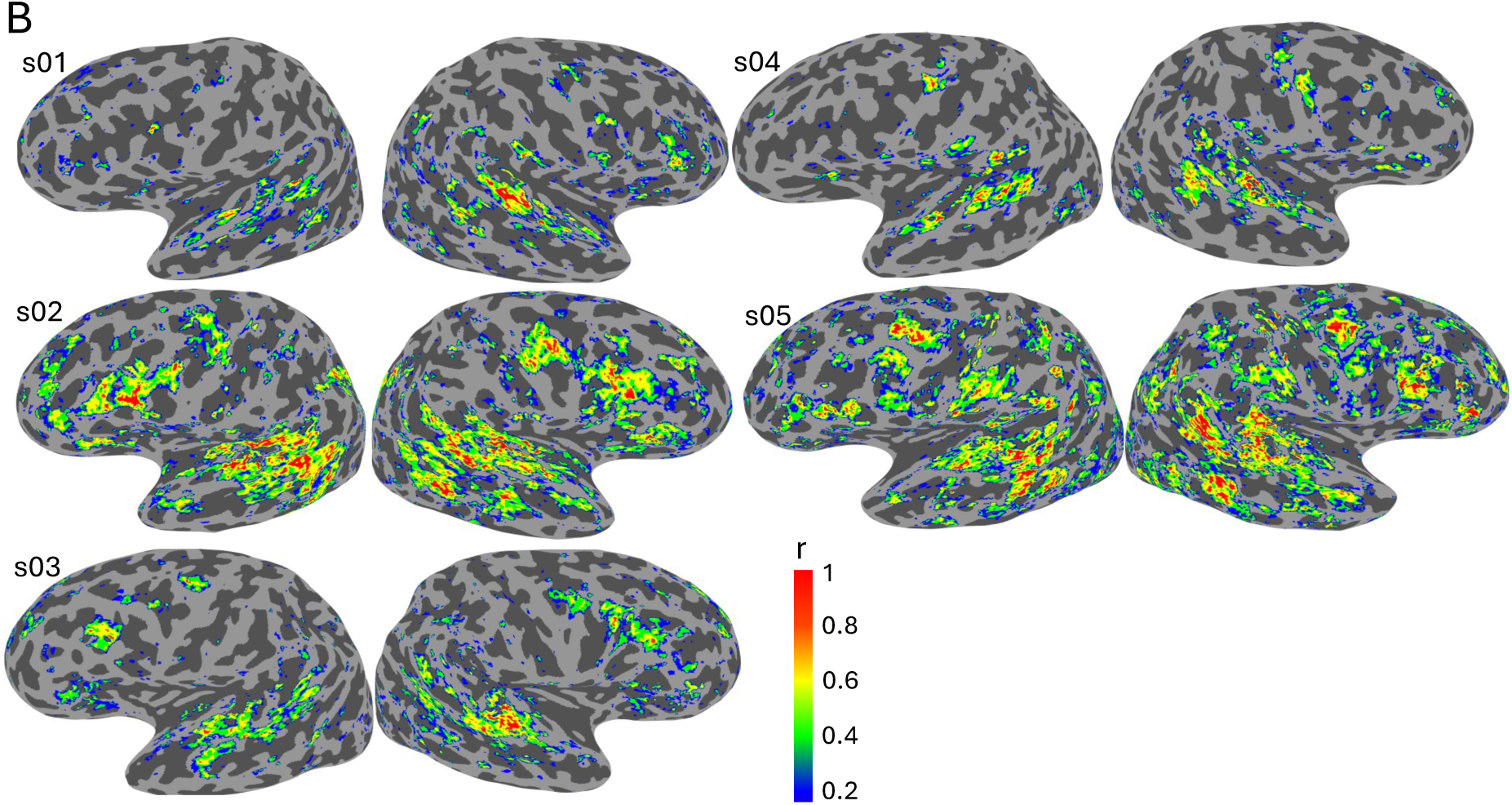
HRF shapes are reproducible across participants and fMRI sessions. **(A)** Each voxel’s cluster group membership is displayed on the cortical surface of each participant. Colours represent cluster groups defined in Figure 3 (far right). **(B)** Each voxel’s response was correlated between Session_1 and Session_2. Response stability across the two sessions was quantified with Pearson correlations and projected onto each participant’s cortical surface. The colour bar represents Pearson correlation coefficient.

Reliable and stereotypical response shapes were observed in auditory temporal regions and surrounding regions across participants (shown in orange and red in Fig. 4A). However, some responses, such as the negative responses from cluster group 3 and 4 (shown in purple and blue in Fig. 4A), were less consistently identified across participants and sessions. Some less typical responses, such as sustained positive responses (shown in yellow, Fig. 4A) were present in all participants and extended beyond auditory temporal regions. These responses were more pronounced in some participants compared with others (e.g., s02).

Response patterns were generally stable across the two sessions, as quantified by correlating the voxelwise event-related averages between Session_1 and Session_2 (median Pearson’s correlation across all voxels and participants: 0.365, SD = 0.349). However, stability varied across brain regions as can be seen in the Pearson correlation coefficients displayed on participants’ cortices (Fig. 4B). Shapes more similar to the canonical HRF (e.g., group 6 and 7) were highly consistent across sessions and exhibited higher correlation coefficients, whereas negative responses (e.g. group 3 and 4) were not as reliably detected across sessions (see Discussion).

#### 3.1.1 Testing custom HRF models against standard approaches in independent data

Within the Original Dataset, based on HRF shapes observed in session 1, six hemodynamically plausible models were created (see 2.3.5 and Fig. 2). These were tested on cluster-level time-series from independent Session_2 data, which were fit to both custom and standard HRF models. (We excluded four clusters that did not result in a reliable fit to any of the six custom models). Overall, the custom models captured significantly more variability of responses within Session_2 data than common HRF models did. After Bonferroni correction, differences between model types remained significant: custom models versus canonical HRF (p <0.001), custom models versus canonical HRF with derivatives (p <0.001), and custom model versus GLM-single basis model (p <0.001).

Examples of responses where standard models did not perform as well as the custom ones are illustrated in Fig. 5. These results suggest that canonical HRF approaches do not optimally capture the diversity of complex response shapes, particularly when the response is fast and narrow (e.g., see canonical model fits for Cluster 21,28 in Fig. 5).

**Fig. 5.**
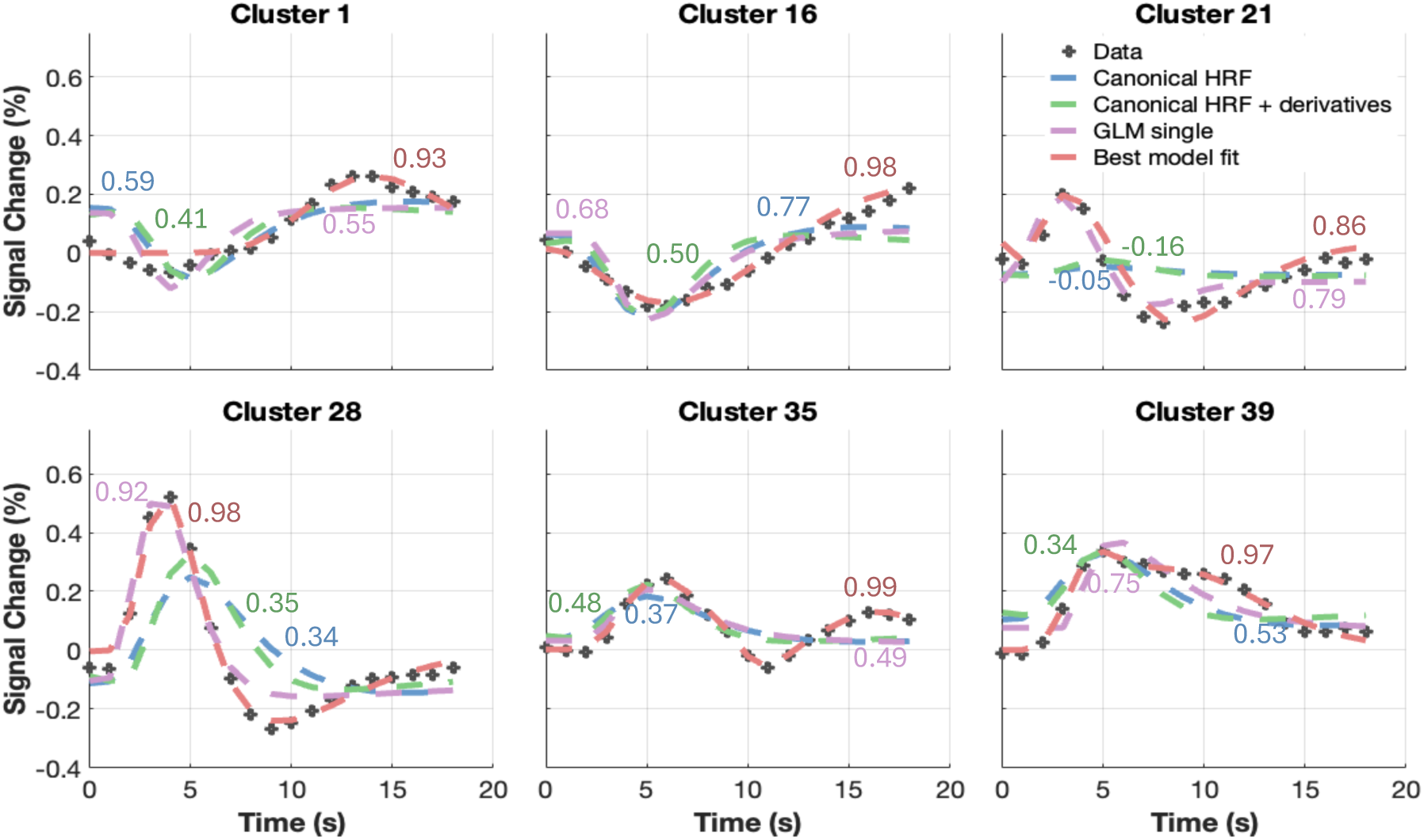
Standard HRF models could not optimally capture observed responses. The displayed cluster examples illustrate response shapes that are less fully captured by either canonical HRF or GLMsingle basis models. Cluster means derived from independent Session_2 data from the Original dataset were used to assess model performance. After excluding 4 of the 40 clusters because the custom models failed to provide reliable fits, 34 cluster means were evaluated for model accuracy. Cluster means (black crossed lines) were fit with: i) SPM HRF (Canonical HRF; blue), ii) SPM HRF with temporal/spatial derivatives (Canonical HRF + derivatives; green), iii) GLMsingle basis HRFs (best basis function shown in purple), and iv) each of the six data-driven custom models (best-fitting model shown in red). 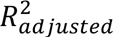 is shown for each fit: canonical HRF (blue), canonical HRF with derivatives (green), GLMsingle basis (purple) and best-fitting data-driven model (red), illustrating relative goodness-of-fit across approaches.

### 3.2 Hemodynamic responses reach stability early and stay consistent over time

#### 3.2.1 Relationship between data amount and goodness of fit

After averaging voxel responses over different run combinations, we found that – unsurprisingly - including more data increased the goodness of fit (measured via 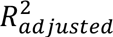) of the best fitting model, for both simulated (see Methods 2.3.6) and the observed data for each participant (Fig. 6). The best-fit metric (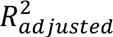) was averaged across voxels and plotted for each run combination, from run 1 to run 6 (Session_1) and up to run 12, representing the average across both sessions (Fig. 5). For the Replication dataset, run 24 corresponded to the average across all four sessions. Across participants, the largest increase in goodness of fit occurred between Run_1 and 2 (Δ0.07 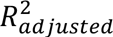 for Original dataset and Δ0.08 for the Replication dataset). Notably, the increase between Run_1 and 2 was smaller within Session_2 in the Original dataset (Δ0.02 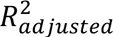) which may reflect fatigue and boredom. Furthermore, in the Original dataset, the average change in goodness-of-fit within Session_1 (Run_1 to 6) was Δ0.2 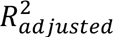. However, when examining changes across sessions in the Original dataset, the improvement from Session_1 to Session_2 (run 6 to 12) was smaller, amounting to only Δ0.02 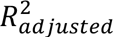. In summary, goodness-of-fit steadily improves as more data are added, with larger gains occurring in earlier compared with later runs (Fig. 6).

**Fig. 6.**
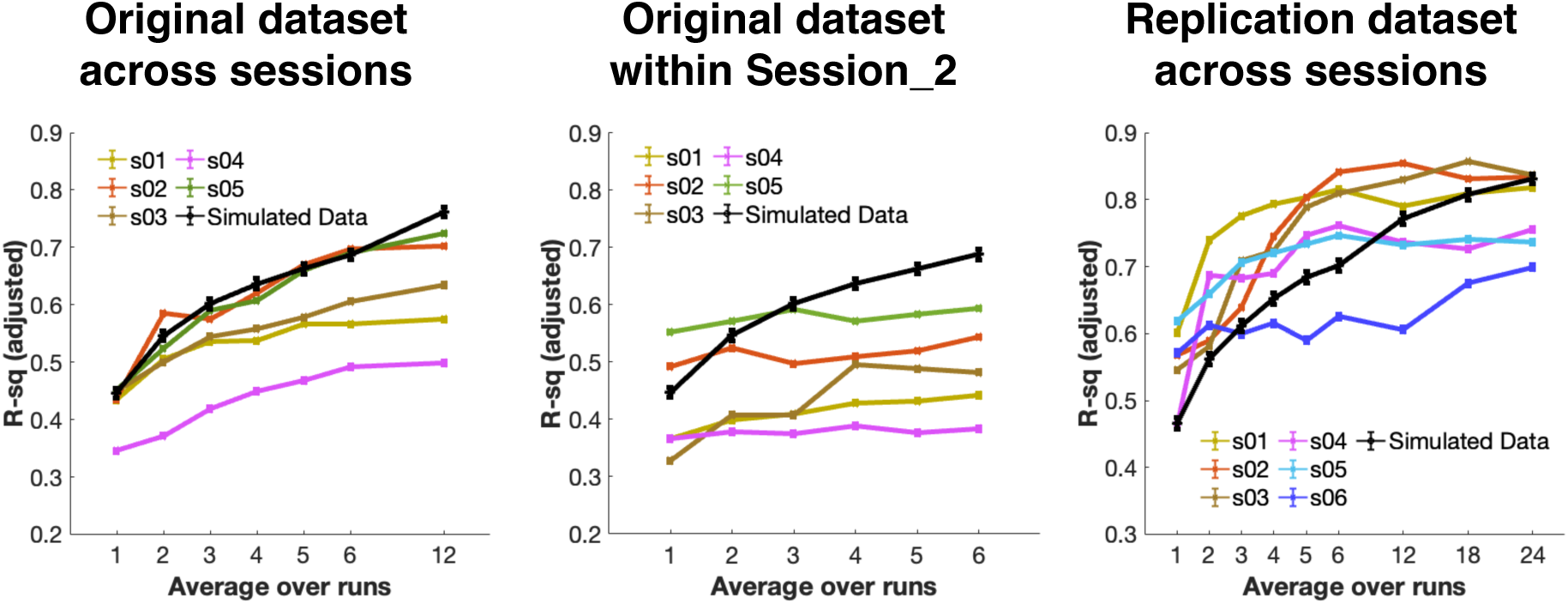
HRF shape estimates (R^2^_adj_) converge within a few runs. The highest 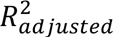 (i.e., the best fitting model for each voxel) were averaged across voxels and plotted for simulated data (black line) and observed participant data (coloured lines). The x axis corresponds to cumulative averages across run combinations (e.g., run 1; runs 1–2; runs 1–3). Adding data led to improvements in model fit of the simulated data. Similarly, in session 1 of the observed data, averaging over larger numbers of trials produced progressively better fits across participants. In Session_2, the fit does not improve with additional runs, contrary to predictions from the simulated data. For the Replication dataset, fit quality increased from Session_1 to Session_2, with a smaller additional improvement through session4. Participants 1–4 are common to both datasets and are shown in consistent colours; other participants are shown in distinct colours. Error bars represent the standard error across voxels.

#### 3.2.2 Spatial representation of stability

Figure 7 illustrates the spatial distribution of this stability on the cortical surface, indicating that stable HRF estimates can be obtained relatively early in a scanning session across participants, especially in auditory temporal regions. On average, across all participants and voxels, 80% of the maximum fit quality was reached within 4.4 runs, suggesting approximately 4 runs are sufficient to obtain stable HRF estimates. However, Fig. 7 shows that stability is reached even earlier in auditory temporal regions, further supporting the high reliability of responses in these regions (see Fig. 4A).

**Fig. 7.**
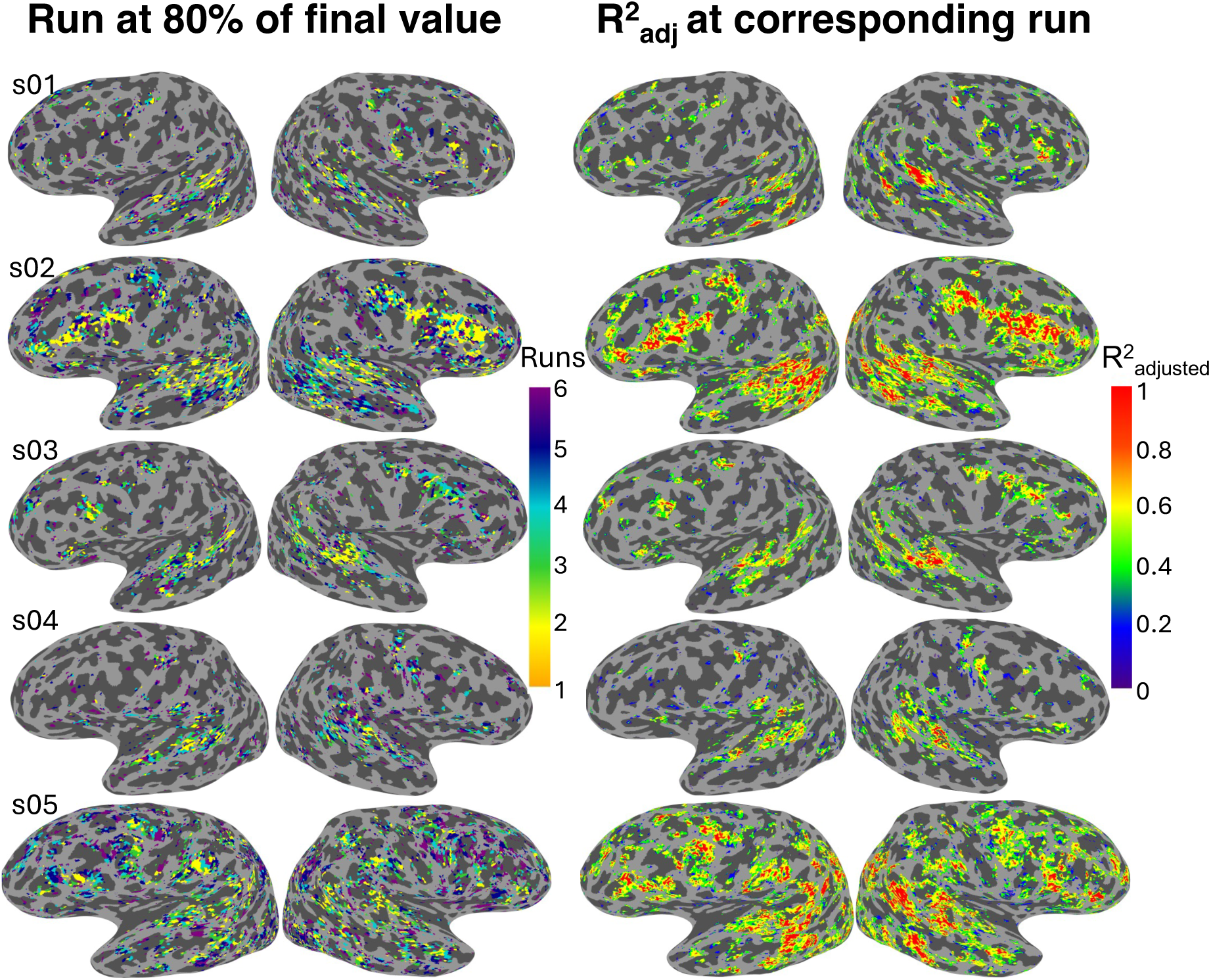
HRF shapes stabilise rapidly within a session in auditory regions. The run at which each voxel’s 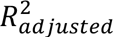 reached 80% of the final value (average across six runs) is shown in each participant’s cortical surface (left), along with the associated 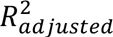 at that run (right). Colour bars represent run number (left) and 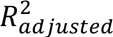 (right) respectively. Data is shown for Original dataset Session_1.

### 3.3 Response characteristics in auditory regions follow reproducible spatial patterns

Given the relatively high stability of responses in auditory temporal regions (Fig. 4B & 7), we next investigated whether systematic differences in peak latency or amplitude could be detected within these regions. Values were averaged across hemispheres given the absence of *a priori* hypotheses regarding hemispheric differences in peak latency or amplitude and were z-score normalised within each participant to control for inter-individual differences by subtracting the mean across that participant’s ROIs and dividing by the corresponding standard deviation. These Z-score values were averaged across participants to generate peak latency and amplitude maps. Despite inter-individual differences (reflected in individually tailored colourbars, see supplementary Fig. 1 and axes in Fig. 9), consistent general patterns in peak latency and amplitude were evident across participants (Fig. 8). More specifically, faster responses were localised to medial regions (particularly in the putative primary auditory ROI “A1/R”, shown in medium blue), whereas slower responses (orange-red) localised to lateral regions (e.g., putative secondary auditory ROIs like “RA4”).

**Fig 8.**
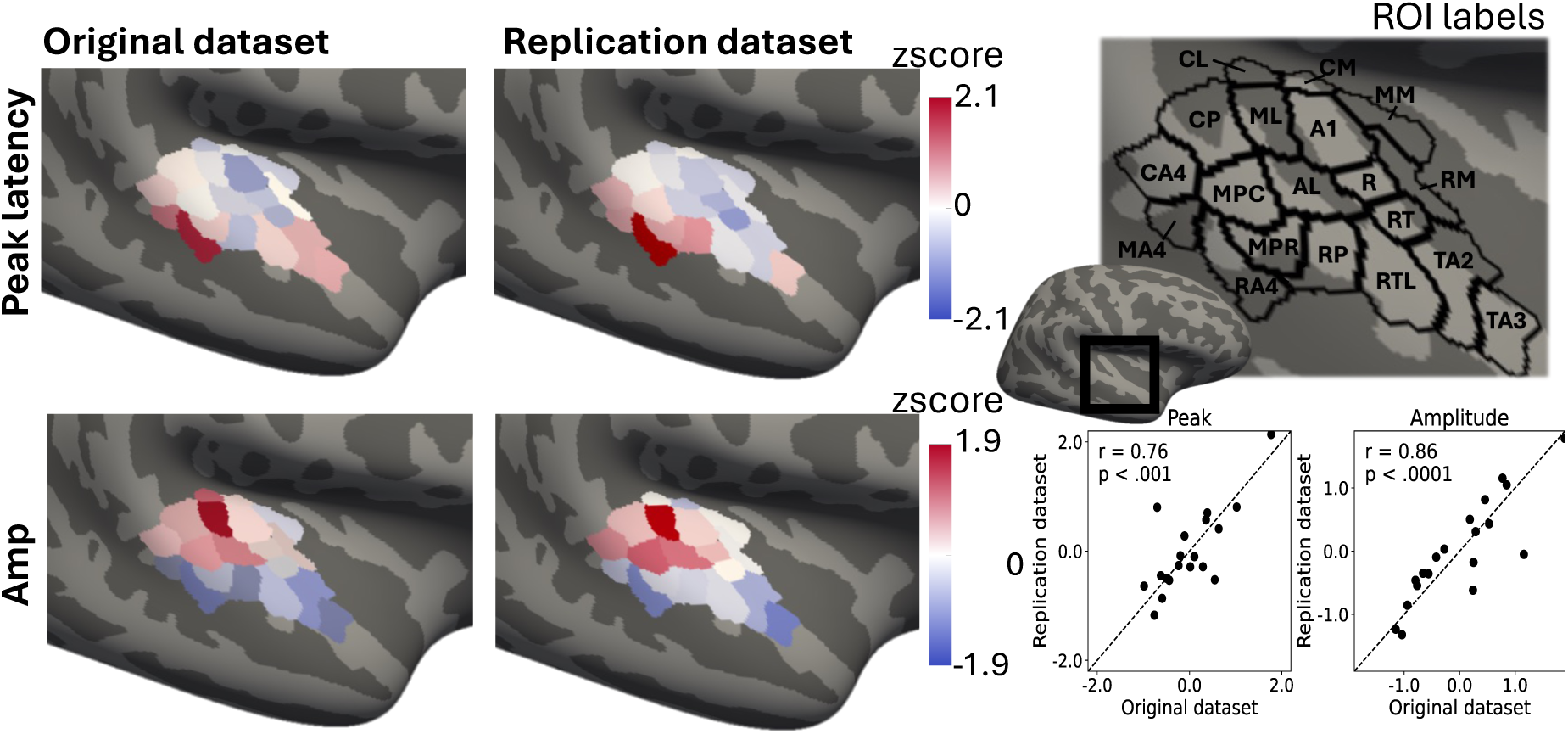
Peak latency and amplitude vary systematically across auditory regions. Peak latency values were normalized within each participant by converting them to z-scores. For visualisation, these z-scores were averaged across participants for each ROI and dataset and then mapped to a colour scale. ROIs are shown on the fsaverage cortical surface. A symmetric colour scale was used, with limits held constant (±2.1 for peak and (±1.9 for amplitude) across datasets. N=5 for Original dataset, N=6 for Replication dataset. See Supplementary Table 2 for ROI annotations.

**Fig. 9.**
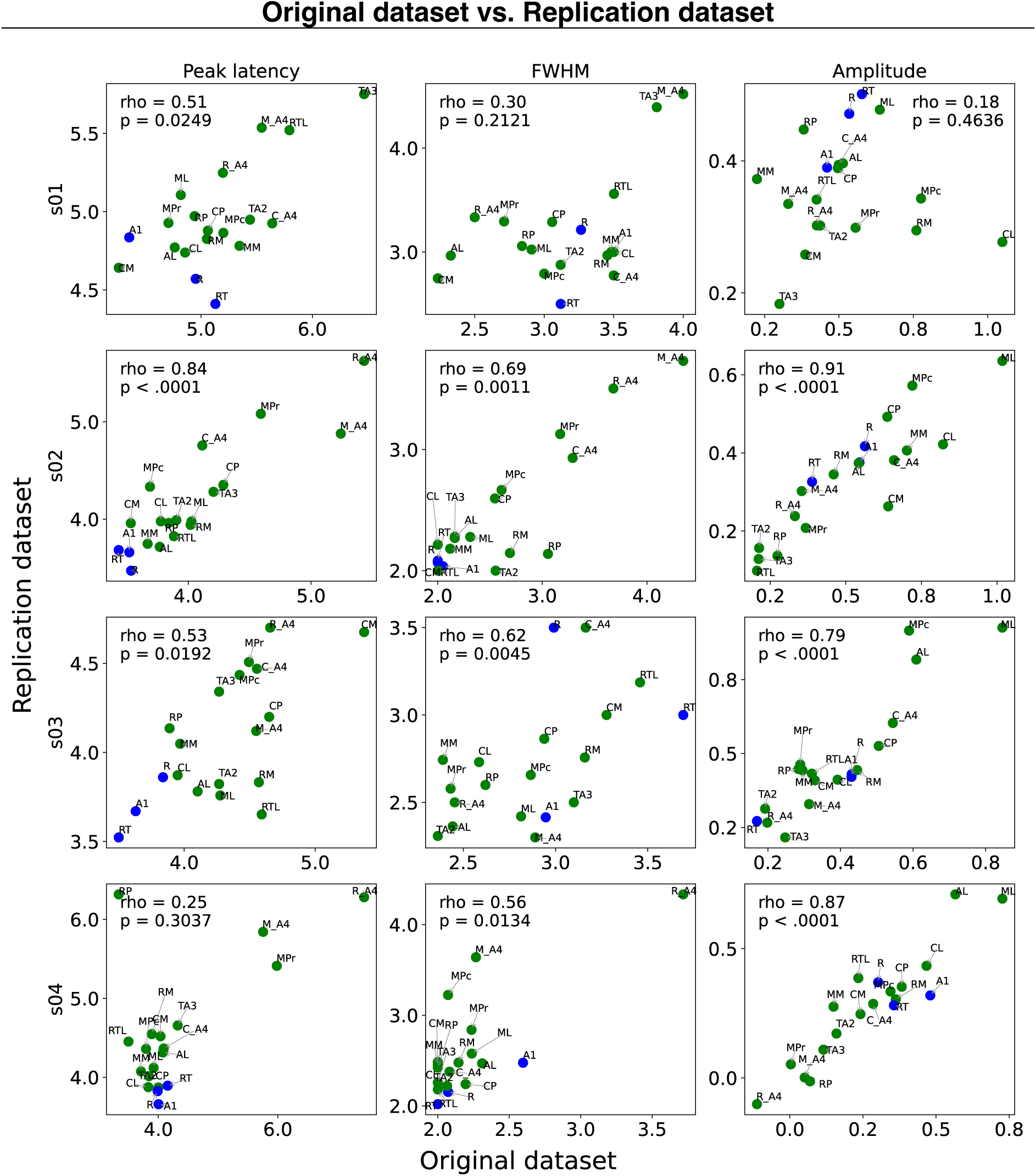
Temporal characteristics of the HRF are reproducible across datasets. Peak latency (s), FWHM (s) and amplitude (% signal change) of the best-fitting model per ROI were averaged across hemispheres and correlated across the two independent datasets for each participant. For Participants 2–4, values showed positive correlations between the two independent acquisitions, with the strongest correspondence observed for amplitude. In contrast, for Participant 1, only peak latency exhibited a significant positive correlation between datasets, whereas width and amplitude showed weaker correspondence. Overall, for three out of four participants HRF characteristics were largely reproducible across auditory ROIs in the two independent datasets. Original dataset values are shown on the x-axis and Replication dataset values are displayed on the y-axis. Spearman’s rho and corresponding p-values are shown for each parameter-participant combination.

Correspondingly, regions with faster responses also tended to show higher response amplitudes, reflecting a coupled HRF peak latency and amplitude gradient, as supported by additional analyses (see supplementary Fig. 2 for the negative relationship between peak latency and amplitude). Crucially, very similar spatial patterns were observed in the Replication dataset collected with a different protocol which included both participants from the first dataset and additional new participants, supporting the reliability of these effects (see Fig. 8 and 9 for cross-dataset correlations). More specifically, Pearson’s correlations revealed both strong correlations for peak latency (r= 0.76, p <.002) and amplitude (r=0.86, p<.0001) between the two datasets.

To further assess reproducibility across acquisitions, we correlated the HRF temporal characteristics using Spearman correlation (peak latency, width, and amplitude) between the Original and Replication datasets for participants common to both datasets. These parameters showed similar relationships across the two datasets, with peak latency and width being positively correlated, whereas peak latency and amplitude showed a moderate negative relationship (see Supplementary Figure 2). Figure 9 displays correlations for each participant, with values based on the best-fitting model for each ROI that were averaged across hemispheres.

Putative primary auditory ROIs “A1”, “R” and “RT”, which were consistently among the fastest ROIs for both datasets (Fig. 8, based on z-scores), are highlighted in blue in Fig. 9 to facilitate tracking of their HRF characteristics in each participant. Though more variable, amplitude in these ROIs was in the mid-to-high ranges (highlighted in medium red in Fig. 8) compared to lateral regions (blue, Fig.8) and FWHM varied more across participants (Fig. 9). The faster and stronger magnitude responses observed here are in line with work from the electrophysiological and intracranial literature, which suggests that these regions exhibit sharper responses compared to surrounding regions (Benson & Hienz, 1978; Liégeois-Chauvel et al., 1994; López-Madrona et al., 2024).

### 3.4 Relationship between HRF properties and underlying cortical properties

Given the known myelination profiles of the ROIs with the fastest observed peak times in the current study (i.e., A1, R, RT), we investigated whether cortical R_1_ values – strongly driven by proportion of myelin content (Lutti et al., 2014; Mottershead et al., 2003; Schmierer et al., 2008) could explain variability in peak latency and amplitude across ROIs. Voxelwise responses were averaged per ROI and then fit to the data-driven models. In a linear mixed-effects model peak latency or amplitude were specified as a dependent variables, whereas the predictor was R_1_. Hemisphere was included as a fixed effect but had no significant effect on the outcome variable so data are shown pooled across hemispheres. Analysis of the Original dataset revealed that peak timing could be predicted by R_1_ values within auditory ROIs, indicating that regions with higher degrees of myelination exhibited faster peak times (β = −7.92, p =0.0024). Although similar in slope, the relationship between R_1_ and peak timing was not significant in the Replication dataset (β = −4.76, p = 0.0802). Amplitude showed a significantly positive relationship with myelination in both the Original and the Replication dataset (Fig. 10), where higher magnitude of responses in auditory regions corresponded to higher degrees of putative myelination.

**Fig. 10.**
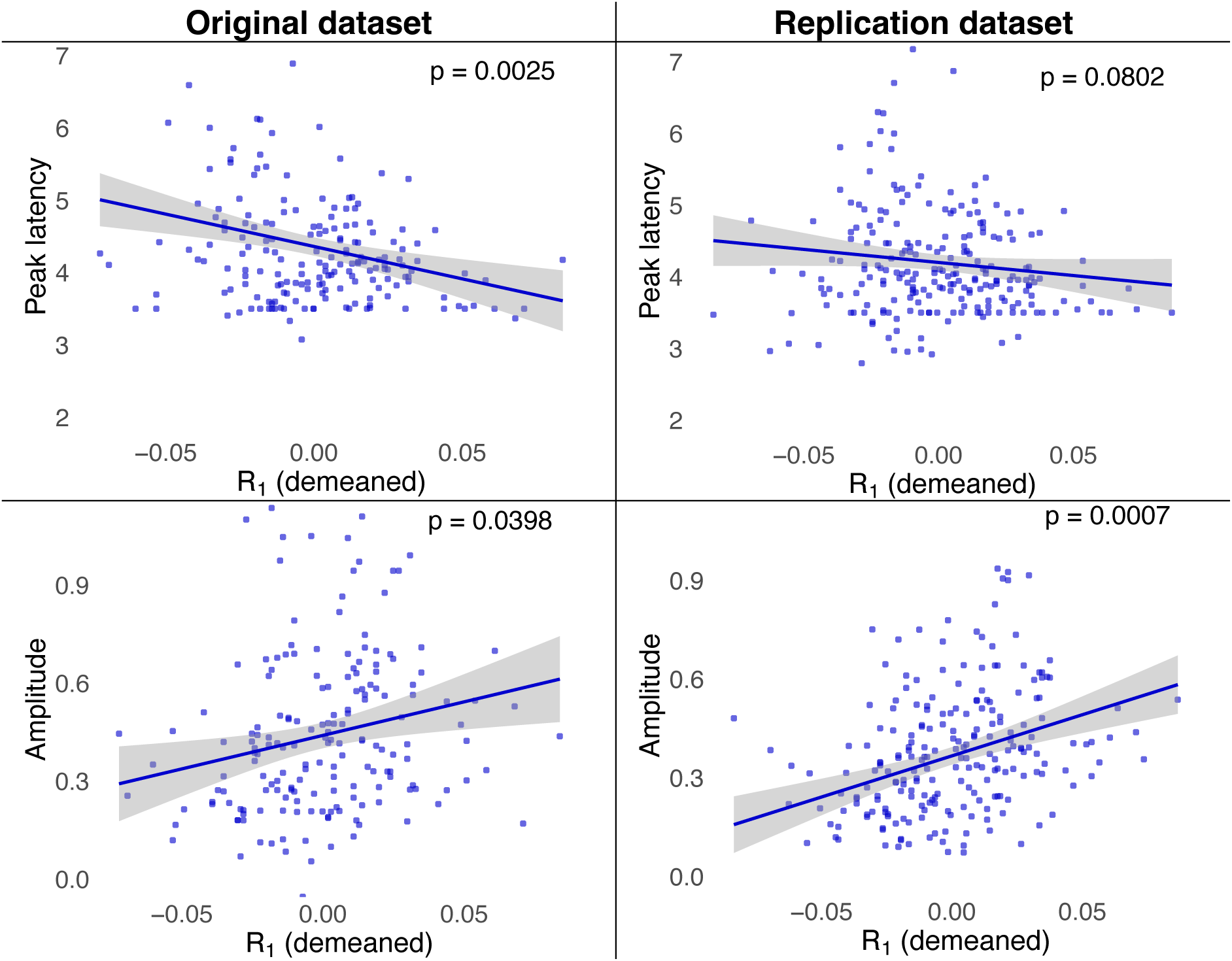
HRF peak latency and amplitude can be predicted by R_1_ values in auditory regions. The relationship between R_1_ and HRF peak latency, and R_1_ and peak amplitude, was estimated using linear mixed-effects models and is shown here for both datasets. The results show a negative relationship between R_1_ and peak latency and a positive relationship between R_1_ and amplitude, where ROIs with higher R_1_ values tend to show faster peak onsets, and higher peak amplitude at the time of peak. Each point represents an ROI-level measurement (e.g., peak latency vs demeaned R_1_), with R_1_ values demeaned across participants to facilitate interpretation of the intercept (0 = average across participants). The solid line shows the linear fit and the shaded region represents the 95% confidence interval. Hemisphere was included as a fixed effect in the model but is not shown in the plot, as it had no significant effect on the outcome variables. The p-value is indicated on the figure.

## 4 Discussion

### Summary of findings

Using ‘fast fMRI’ to characterise voxelwise responses in the brain when participants listen to short meaningful sounds, we found stable and complex responses that were reproducible across participants and sessions. Many of the observed responses would not be fully captured using canonical HRF models, even when including temporal derivatives. Having acquired up to 24 runs per person, we confirmed that accumulating data was associated with an improved ability to capture responses reliably. However, from a practical standpoint, four approximately 4-minute runs were typically sufficient to get a stable estimate of the HRF across the brain. Even fewer runs were necessary for estimates to converge in auditory temporal regions. Within these auditory regions, we found that temporal characteristics of the hemodynamic response exhibited relatively similar spatial profiles across participants. Notably, we observed faster and higher magnitude responses in medial auditory regions and slower and lower magnitude responses in more lateral auditory regions. Lastly, we found that R_1_ values predicted peak latency (in one dataset) and amplitude (in two datasets) in auditory regions, consistent with faster peaks and higher amplitudes in regions with higher degrees of myelination.

### Evidence for non-traditional HRF shapes

The present findings underscore the importance of adopting flexible modelling approaches when analysing timecourses at the voxel or small ROI level, particularly given that poor fits between observed response shapes and standard HRF models can inherently compromise statistical inference. We find that peak times of many of the observed responses were faster than would be expected from canonical models. This observation converges with prior work challenging the assumption of linearity, which has reported both larger-than-expected response amplitudes and faster-than-expected responses to short stimuli, and suggests that shorter stimulus durations are associated with increased nonlinearities in the hemodynamic response (Birn & Bandettini, 2005; De Zwart et al., 2009; Hu et al., 2010; Robson et al., 1998; Yeşilyurt et al., 2008). Consistent with prior work, we report a range of one-,two- and three-gamma responses which included positive and negative peaks, aligning with the growing body of literature advocating for increasingly flexible HRF modelling approaches (Gonzalez-Castillo et al., 2012; Handwerker et al., 2004; Lindquist et al., 2009; Taylor et al., 2018). In the context of the current study, we find that these gamma-based models are sufficient and well-suited for capturing auditory hemodynamic responses, which aligns with evidence from motor, somatosensory, and visual cortices (Bailes et al., 2023; De Zwart et al., 2005a; Siero et al., 2011). A key advantage of the models used here is their ability to accommodate a broad range of response shapes beyond the two-gamma shapes, while remaining straightforward to implement. At the same time, GLMsingle performed well overall and represents a viable alternative for flexible HRF modelling, although its ability was somewhat limited in capturing responses with pronounced late negative peaks following delayed positive peaks, or responses comprising more than two distinct peaks.

### The potential of fast fMRI in auditory research and methodological challenges

Using faster TRs at 3T, we characterised the auditory hemodynamic response function with greater temporal precision than typically reported in previous fMRI studies (De Zwart et al., 2005; Lewis et al., 2016). We observe quite fine timing differences in the HRF in auditory temporal regions, which complement findings of intra-regional differences in response latency in primary visual regions using high resolution imaging (Gomez et al., 2024). The observed spatial gradient of peak latency is in line with the previously proposed distinction between fast and slow responses in auditory regions from computational modelling (Santoro et al., 2014; Zulfiqar et al., 2021). Intriguingly, these fine-grained timing differences could be predicted by regional variations in R_1_, a proxy for myelination. While we could confirm previous findings that magnitude of the response is linked to R_1_, our results offer initial evidence that HRF timing may also relate to cortical microstructure (Patitucci, 2021). This finding may help shed more light on the relationship between R_1_ and neurovascular coupling and has potential to advance our understanding of auditory processing due to the established role of myelination in the processing of sound (Fletcher et al., 2021). However, further studies are warranted to confirm this result, as the relationship was not significant in the Replication dataset for peak timing, despite being qualitatively similar. Further studies with larger sample sizes will be required to establish the robustness of this effect. By comparison, the positive relationship between R_1_ and initial peak amplitude was more robust, again demonstrating a relationship between BOLD dynamics and underlying macromolecular content (see also Dick et al., 2017).

Direct comparisons between our findings and neural evidence need to be performed with caution, since latency differences in HRF can arise from a multitude of factors, including vascular differences. If they are related to neural activity at all, they are likely to be due to the duration rather than onset of underlying neural activity (Henson et al., 2002). Although spatial resolution in fMRI still limits direct comparisons with invasive animal work, we suggest that the current work represents a useful step towards bridging this gap by overcoming a range of challenges.

Critically, we aimed to obtain greater spatial specificity by using voxelwise or small ROI-wise HRFs rather than averaging across large regions, which can obscure fine-grained differences in HRF timing and amplitude. Second, using relatively high spatial resolution (2 mm isotropic) for 3T, and whole-brain imaging, we captured much of the auditory response across brain regions while maintaining reasonable spatial resolution. The 2 mm isotropic resolution also lessens macrovascular contributions at 3T compared to larger voxel sizes (e.g., 3×3×3 mm) which disproportionately capture signals from large veins near the cortical surface, producing large BOLD signals far from the underlying neural activity (Triantafyllou et al., 2005) and biasing signal amplitude towards large veins on the cortical surface (Moerel et al., 2018). To mitigate this, we excluded low TSNR voxels and applied surface-based smoothing (2 mm FWHM). This correction is particularly important in auditory cortex due to its unique vascular architecture and proximity to large diving veins, which can bias BOLD signal away from underlying neural activity (Faes et al., 2023; Moerel et al., 2018; Taylor et al., 2022). While acknowledging that significant partial volume effects are still present even at high spatial resolution, it is unlikely that responses observed in auditory regions are driven to any significant extent by microvasculature due to the reliable fast peaks and relatively low amplitudes. It is also worth noting that the use of higher temporal resolution (TR=1000ms) allowed us to examine earlier phases of the response with more detail, such as the initial dip, which have been previously linked to neural processing (Polimeni & Lewis, 2021; Zaidi et al., 2023).

### Limitations and outlook significance statement

The work presented here comes with some limitations. We focused on BOLD responses to fast, temporally spaced stimuli, to characterize the full hemodynamic response. Because the design employed a passive listening task, it is difficult to separate neural from neurovascular contributions to differences in the response. Even though participants were monitored and instructed to attend to the stimuli, no behavioural feedback was collected to confirm engagement, and the passive nature of the task may have increased boredom or habituation, potentially contributing to reduced response amplitudes in later sessions. It is notable that negative responses (e.g., Groups 3 and 4 in 3.1) were not as reliably detected across sessions, indicating that these slower negative components may reflect a combination of genuine neural variability across trials and sessions as well as hemodynamic factors. Future studies could apply the data-driven models developed here to study the hemodynamic response to different tasks in the same participants to help disentangle neural from vascular effects potentially in combination with MEG or, potentially simultaneously-acquired fMRI and EEG.

In order to investigate the stability of the HRF over time, we prioritized acquisition of many datapoints in the same expert participants above obtaining a larger-N sample with fewer runs and sessions. While benefitting from the novel cross-session registration techniques that allowed us to compare voxelwise responses over different sessions in the same participant, this choice necessarily limits the potential for group-wide for broad generalisation to the population level, as illustrated by the negative responses that were largely driven by one participant and not as stable across sessions as more canonical responses. Thus, future studies on the HRF variability at higher resolution would benefit from larger sample sizes to confirm whether such patterns can be observed in a representative sample.

It is also worth noting that high temporal resolution approaches are affected by lower image SNR and while effects can be demonstrated at the individual level, they often don’t persist at the group level (Kirilina et al., 2016; Zeidman et al., 2019). This is exemplified by the observation of the initial dip response in the current work. While the higher temporal resolution allowed us to get a more detailed insight into the early stages of the response and individual responses exhibited an initial dip – none of the determined clusters included an initial dip. Even when looking at participant-wise responses, averaging over several voxels already eliminated the initial dip response which has been reported before (Watanabe et al., 2013). Detecting the initial dip is further compromised because of its small amplitude and high variability compared to later time points (e.g., the main positive peak) of the response. Additionally, modelling the dip carries a substantial risk of false positives, as noise can easily be misinterpreted as a genuine dip. This underscores a key limitation of our approach—the use of flexible and complex models, which increases susceptibility to overfitting. We tried to minimize the effects of overfitting as much as possible by adjusting the goodness of fit measures according to the number of parameters of the models and fitting the models on independent data for validation. In spite of this, the high goodness of fit measures of our data-driven models may in part arise from overfitting and thus the comparison of our flexible models to more rigid canonical models need to be viewed critically.

## Conclusion

Our findings demonstrate that auditory HRFs are diverse, reproducible, and regionally specific. By moving beyond canonical models and leveraging fast fMRI and multimodal imaging, we provide a framework for more accurate and biologically meaningful characterisation of auditory processing in the human brain. This framework facilitates closer links with electrophysiology to improve understanding of auditory temporal dynamics and highlights the need for updated HRF modelling in auditory neuroscience.

## 5 Data and Code Availability

The data used in this paper has been deposited on OpenNeuro: doi:10.18112/openneuro.ds007780.v1.0.0

Preprocessing and analysis scripts used for this paper are documented here: https://github.com/LETSCHN/auditory-hrf

## 6 Author contributions

Conceptualisation: L.S., I.Z., F.D.; Methodology: L.S., I.Z., Y.B., F.D.; Software: I.Z., Y.B.; Validation: L.S., I.Z., F.D.; Formal Analysis: L.S., I.Z., Y.B., F.D.; Investigation: L.S., I.Z., F.D.; Resources: L.L.H., F.D.; Data Curation: L.S.; Writing - Original Draft: L.S., I.Z., F.D.; Writing - Review and Editing: L.S.,I.Z.,Y.B.,L.L.H.,M.F.C.,F.D.; Visualisation: L.S.,I.Z.,Y.B.; Supervision: I.Z.,F.D.; Project Administration: L.S., I.Z., F.D.; Funding Acquisition: L.L.H.,F.D.

## 8 Supplementary Material

### Appendix 1 Cross-session registration

Our implementation of groupwise registration leverages tools implemented in the nitorch toolbox (https://github.com/balbasty/nitorch). A script that contains all necessary steps is available in the nitorch repository (https://github.com/balbasty/nitorch/blob/master/scripts/optinorm.bash). Each pair of images was co-registered by optimising their local correlation coefficient. The deformation model used in this routine combines an affine transformation, encoded by its Lie algebra (Ashburner & Ridgway, 2013), and a nonlinear displacement field, encoded by a stationary velocity field (SVF) (Ashburner, 2007). Importantly, the affine operates on both sides of the nonlinear transformation, ensuring that it is located in the average space of both images. Formally, it means that the transformation from session A to session B can be written as 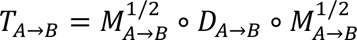, where ∘ denotes composition; *X*^1/2.^ denotes the matrix square root of *X*; *M_A→B_* is the affine transformation; and *D_A→B_* = exp (*V_A→B_*) is the nonlinear transformation exponentiated from its stationary velocity field *V_A→B_*. The transformation from session A to the average space of sessions A and B can be written as 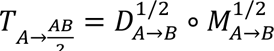, where 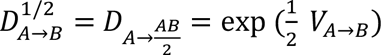, and the transformation from session B to the average space of sessions A and B can be written as 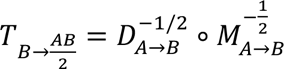, where 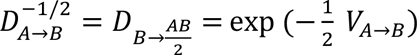, The registration routine outputs the affine matrix *M_A→B_*, the SVF *V_A→B_*, and the Hessian of the SVF *H_A→B_*. Once done, we are left with a set of transformations *T_Ai→Aj_*. One can consider each space *A*_!_ as a node in a graph, and each transformation *T_Ai→Aj_* as an edge (Casamitjana et al., 2025; Iglesias et al., 2018; Leung et al., 2012). An average node *A* can be inserted, along with the unknown edges 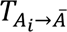. The values of the missing edges are obtained by minimising the squared distance between each known edge *T_Ai→Aj_* and the equivalent unknown path 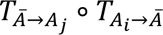 Our implementation differs from existing approaches in two ways:

1. When computing the affine average space, we use a nonlinear optimisation rather than linearising the problem.
2. When computing the nonlinear average space, we do linearise the problem but weight the squared distance by the Hessian of the SVF, which ensures that errors are preferentially distributed across non-informative voxels.

Finally, we obtain a set of optimal transformations 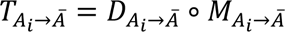, which are used to reslice each session into their optimal average space.

**Supplementary Table 1.**
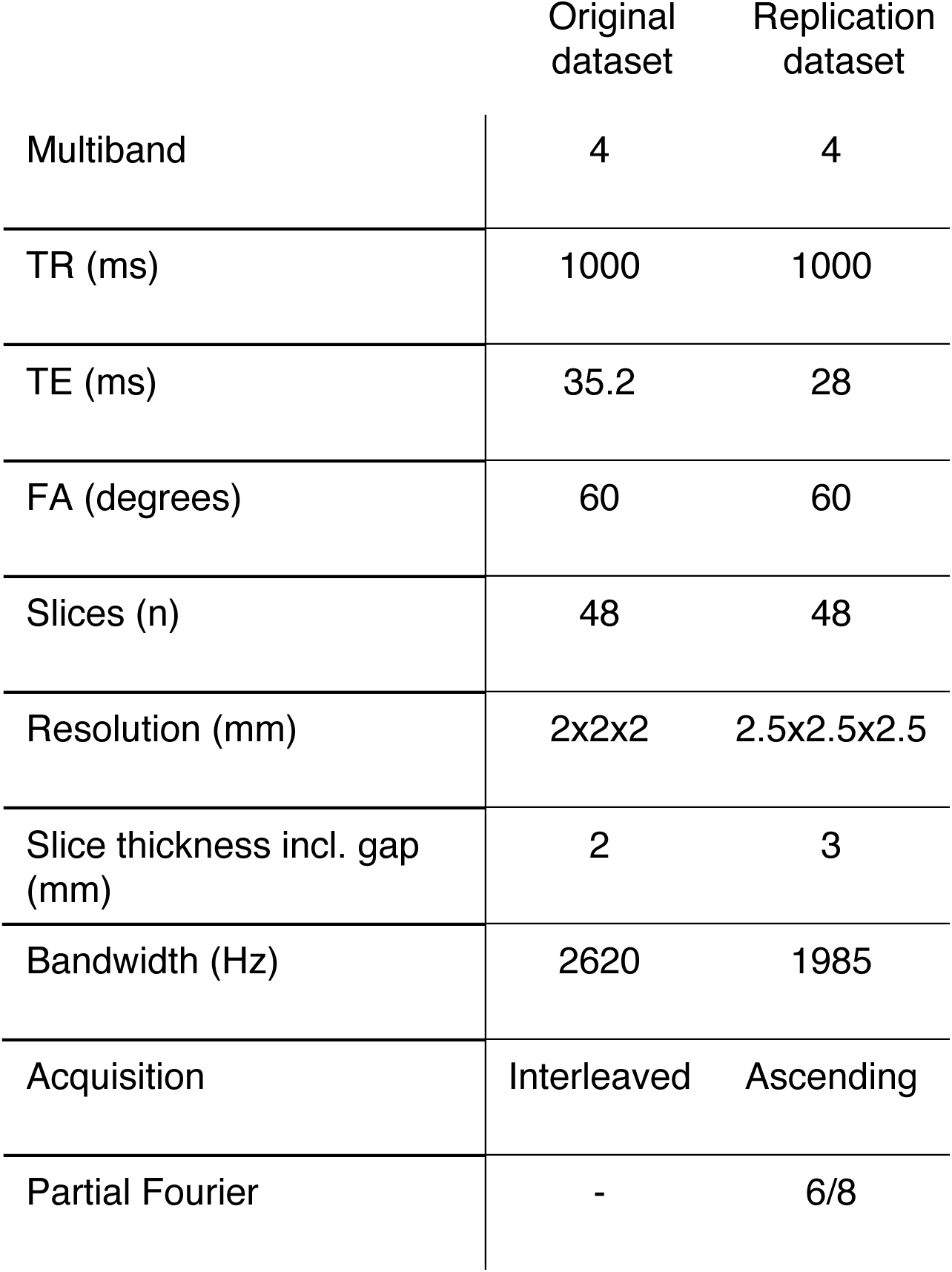
Overview of two acquisitions (Original dataset and Replication dataset)

| Original<br>dataset | Replication<br>dataset |
| --- | --- |

|  |  |  |
| --- | --- | --- |
| Multiband | 4 | 4 |
| TR (ms) | 1000 | 1000 |
| TE (ms) | 35.2 | 28 |
| FA (degrees) | 60 | 60 |
| Slices (n) | 48 | 48 |
| Resolution (mm) | 2x2x2 | 2.5x2.5x2.5 |
| Slice thickness incl. gap (mm) | 2 | 3 |
| Bandwidth (Hz) | 2620 | 1985 |
| Acquisition | Interleaved | Ascending |
| Partial Fourier | - | 6/8 |

**Supplementary Fig. 1.**
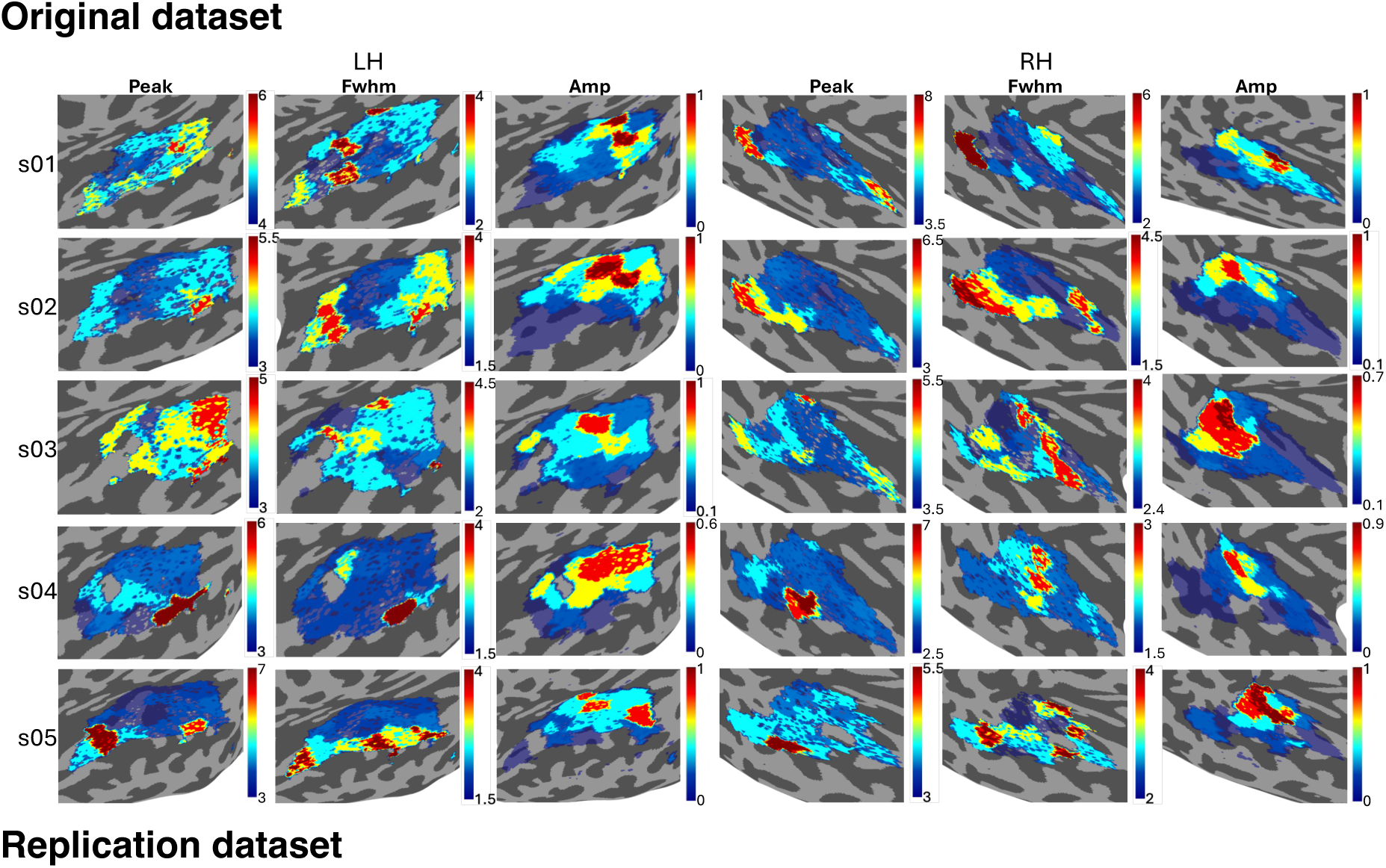
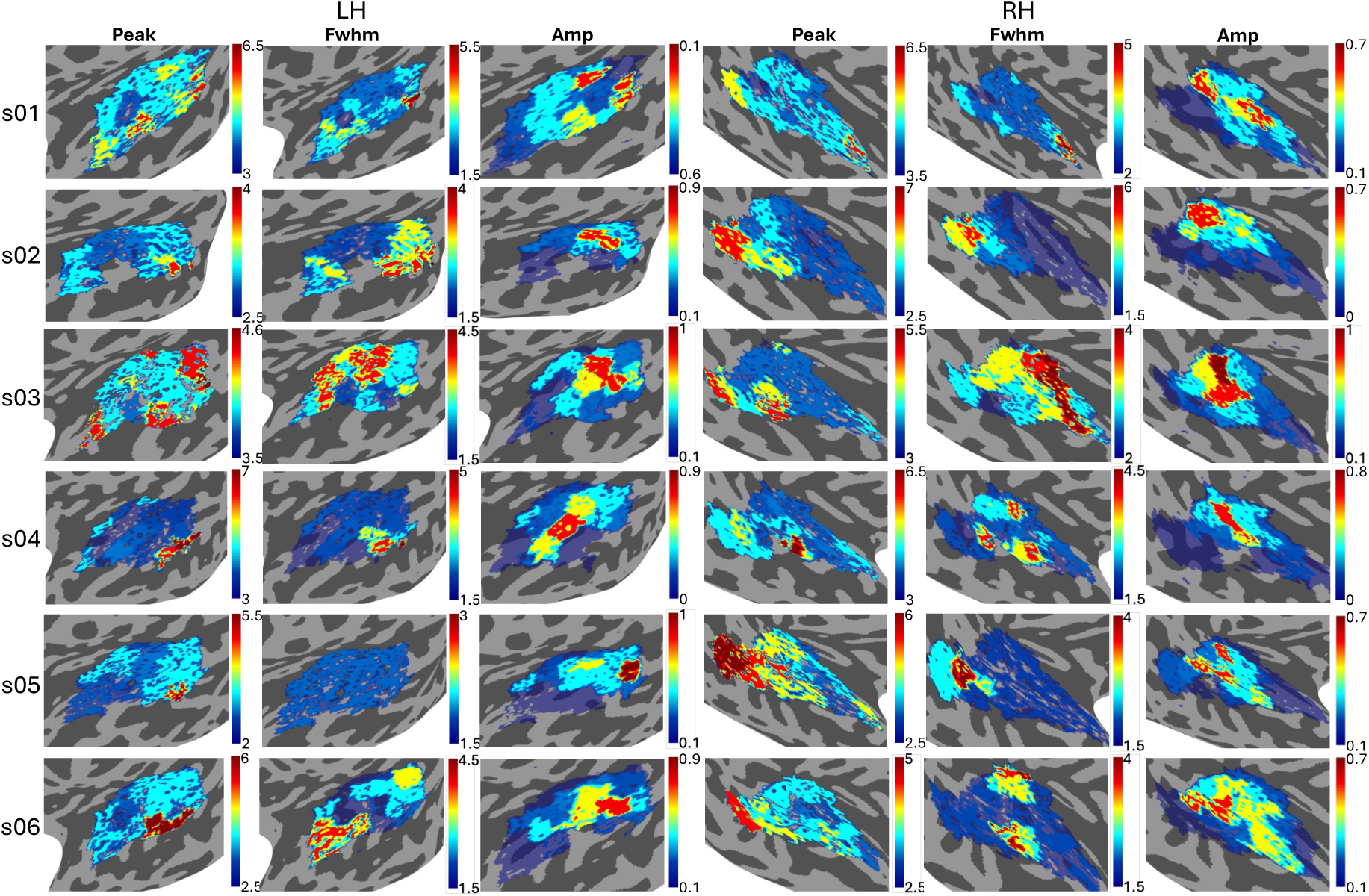
Responses across auditory brain regions exhibit similar characteristics across participants. **(A)** Peak (s), FWHM (s) and amplitude (% signal change) of the first peak of the best model are shown for temporal auditory ROIs on each participant’s surface. If the best model was either model 5 or 6 which include an initial negative peak (i.e. ‘initial dip’), the second peak was chosen for illustration. ROIs left blank either had poor fits (<0.2 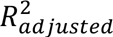) or were very small (<5voxels). Participants 1-4 are identical for both datasets. Color bars vary per participant to reflect individual ranges in parameters.

**Supplementary Table 2.** Auditory ROI annotations.

|  |  |
| --- | --- |
| *Auditory core (primary)* |  |
| A1 | Primary auditory area |
| R | Rostral auditory area |
| RT | Rostrotemporal area |
| *Auditory caudal/medial belt* |  |
| MM | Middle medial belt |
| RM | Rostromedial belt |
| CM | Caudomedial belt |
| *Auditory lateral belt (secondary)* |  |
| CL | Caudolateral belt |
| ML | Middle lateral belt |
| AL | Anterior lateral belt |
| RTL | Rostrotemporal lateral belt |
| *Auditory para belt (tertiary)* |  |
| CP | Caudal parabelt |
| MPc | Middle parabelt, caudal |
| MPr | Middle parabelt, rostral |
| RP | Rostral parabelt |
| TA2 | Temporal area A, part 2 |
| TA3 | Temporal area A, part 3 |
| *Auditory belt areas, 4th tier (A4)* |  |
| CA4 | Caudal 4th tier auditory |
| MA4 | Middle 4th tier auditory |
| RA4 | Rostral 4th tier auditory |

**Supplementary Fig 2.**
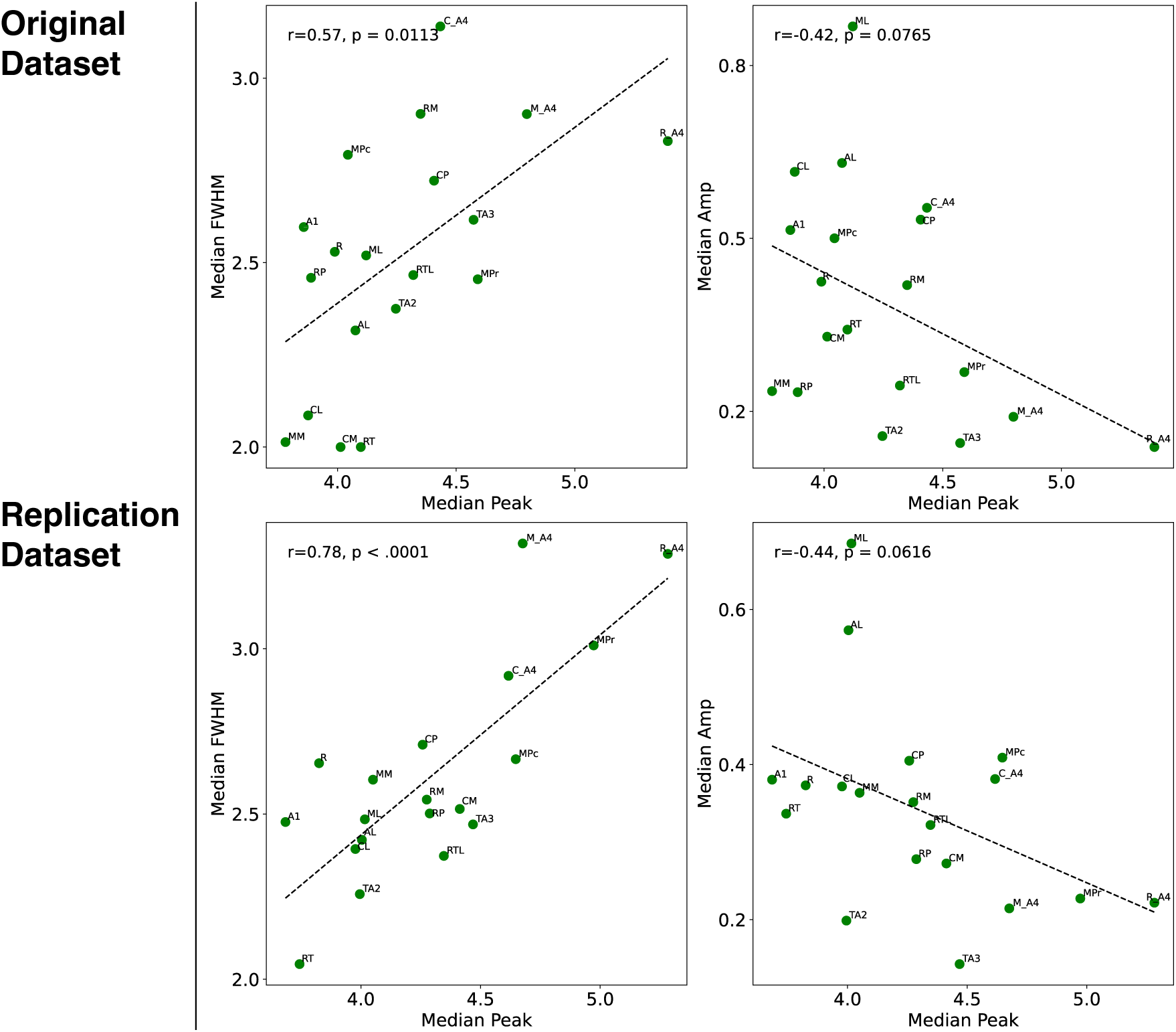
Relationship between time to peak, width and amplitude of responses across datasets. Median of time to peak (s), FWHM (s) and amplitude (% signal change) of the best model per ROI across hemispheres and participants were correlated for each of the two independent datasets. Only data for participants common to both datasets is shown. Peak and width show a strong positive, whereas peak and amplitude show a moderate negative relationship which appears consistent across the two datasets. Pearson correlation coefficient along with p-value are displayed for each comparison.

## Acknowledgements

This work was supported by the National Institutes of Health (R01DC004674), the National Science Foundation (SBE/BCS 2414066, 2420979), the Royal Society Newton Fellowship (NIF\R1\232460) and in-kind support from the Birkbeck/UCL Centre for NeuroImaging (BUCNI).

## References

Ashburner, J. (2007). A fast diffeomorphic image registration algorithm. NeuroImage, 38(1), 95–113. 10.1016/j.neuroimage.2007.07.007

Ashburner, J., & Ridgway, G. R. (2013). Symmetric diffeomorphic modeling of longitudinal structural MRI. Frontiers in Neuroscience, (FEB). 10.3389/fnins.2012.00197

Bailes, S. M., Gomez, D. E. P., Setzer, B., & Lewis, L. D. (2023). Resting-state fMRI signals contain spectral signatures of local hemodynamic response timing. ELife, 12. 10.7554/ELIFE.86453

Baumann, S., Griffiths, T. D., Rees, A., Hunter, D., Sun, L., & Thiele, A. (2010). Characterisation of the BOLD response time course at different levels of the auditory pathway in non-human primates. NeuroImage, 50(3), 1099–1108. 10.1016/j.neuroimage.2009.12.103

Benson, D., & Hienz, R. (1978). Single-unit activity in the auditory cortex of monkeys selectively attending left vs. right ear stimuli. Brain Res., 2(159), 307–320. 10.1016/0006-8993(78)90537-1.

Birn, R. M., & Bandettini, P. A. (2005). The effect of stimulus duty cycle and ‘off’ duration on BOLD response linearity. NeuroImage, 27(1), 70–82. 10.1016/j.neuroimage.2005.03.040

Bollmann, S., & Barth, M. (2021). New acquisition techniques and their prospects for the achievable resolution of fMRI. In Progress in Neurobiology (Vol. 207). Elsevier Ltd. 10.1016/j.pneurobio.2020.101936

Boynton, G. M., Engel, S. A., Glover, G. H., & Heeger, D. J. (1996). Linear Systems Analysis of Functional Magnetic Resonance Imaging in Human V1. In The Journal of Neuroscience (Vol. 16, Number 13).

Buxton, R. B., Uludaǧ, K., Dubowitz, D. J., & Liu, T. T. (2004). Modeling the hemodynamic response to brain activation. NeuroImage, 23(SUPPL. 1). 10.1016/j.neuroimage.2004.07.013

Casamitjana, A., Sala-Llonch, R., Lekadir, K., & Iglesias, J. E. (2025). USLR: An open-source tool for unbiased and smooth longitudinal registration of brain MRI. Medical Image Analysis, 105. 10.1016/j.media.2025.103662

Chen, G., Taylor, P. A., Reynolds, R. C., Leibenluft, E., Pine, D. S., Brotman, M. A., Pagliaccio, D., & Haller, S. P. (2023). BOLD Response is more than just magnitude: Improving detection sensitivity through capturing hemodynamic profiles. NeuroImage, 277. 10.1016/j.neuroimage.2023.120224

Dale, A., & Sereno, M. (1993). Improved localization of cortical activityby combining EEG and MEG with MRI cortical surface reconstruction: a linear approach. J Cogn Neurosci, 5, 162–176.

De Zwart, J. A., Silva, A. C., Van Gelderen, P., Kellman, P., Fukunaga, M., Chu, R., Koretsky, A. P., Frank, J. A., & Duyn, J. H. (2005a). Temporal dynamics of the BOLD fMRI impulse response. NeuroImage, 24(3), 667–677. 10.1016/j.neuroimage.2004.09.013

De Zwart, J. A., Silva, A. C., Van Gelderen, P., Kellman, P., Fukunaga, M., Chu, R., Koretsky, A. P., Frank, J. A., & Duyn, J. H. (2005b). Temporal dynamics of the BOLD fMRI impulse response. NeuroImage, 24(3), 667–677. 10.1016/j.neuroimage.2004.09.013

De Zwart, J. A., van Gelderen, P., Jansma, J. M., Fukunaga, M., Bianciardi, M., & Duyn, J. H. (2009). Hemodynamic nonlinearities affect BOLD fMRI response timing and amplitude. NeuroImage, 47(4), 1649–1658. 10.1016/j.neuroimage.2009.06.001

Dick, F. K., Lehet, M. I., Callaghan, M. F., Keller, T. A., Sereno, M. I., & Holt, L. L. (2017). Extensive tonotopic mapping across auditory cortex is recapitulated by spectrally directed attention and systematically related to cortical myeloarchitecture. Journal of Neuroscience, 37(50), 12187–12201. 10.1523/JNEUROSCI.1436-17.2017

Duvernoy, H. M., Delon, S., & Vannson’, J. L. (1981). Cortical Blood Vessels of the Human Brain. In Brain Research Bulkfin (Vol. 7).

Edwards, L. J., McColgan, P., Helbling, S., Zarkali, A., Vaculčiaková, L., Pine, K. J., Dick, F., & Weiskopf, N. (2023). Quantitative MRI maps of human neocortex explored using cell type-specific gene expression analysis. Cerebral Cortex, 33(9), 5704–5716. 10.1093/cercor/bhac453

Faes, L. K., De Martino, F., & Huber, L. (2023). Cerebral blood volume sensitive layer-fMRI in the human auditory cortex at 7T: Challenges and capabilities. PLoS ONE, 18(2 February). 10.1371/journal.pone.0280855

Feinberg, D. A., Moeller, S., Smith, S. M., Auerbach, E., Ramanna, S., Glasser, M. F., Miller, K. L., Ugurbil, K., & Yacoub, E. (2010). Multiplexed echo planar imaging for sub-second whole brain fmri and fast diffusion imaging. PLoS ONE, 5(12). 10.1371/journal.pone.0015710

Fletcher, J. L., Makowiecki, K., Cullen, C. L., & Young, K. M. (2021). Oligodendrogenesis and myelination regulate cortical development, plasticity and circuit function. In Seminars in Cell and Developmental Biology (Vol. 118, pp. 14–23). Elsevier Ltd. 10.1016/j.semcdb.2021.03.017

Friston, K. J., Fletcher, P., Josephs, O., Holmes, A., Rugg, M. D., & Turner, R. (1998). Event-Related fMRI: Characterizing Differential Responses.

Glasser, M. F., Coalson, T. S., Robinson, E. C., Hacker, C. D., Harwell, J., Yacoub, E., Ugurbil, K., Andersson, J., Beckmann, C. F., Jenkinson, M., Smith, S. M., & Van Essen, D. C. (2016). A multi-modal parcellation of human cerebral cortex. Nature, 536(7615), 171–178. 10.1038/nature18933

Gomez, D. E. P., Polimeni, J. R., & Lewis, L. D. (2024). The temporal specificity of BOLD fMRI is systematically related to anatomical and vascular features of the human brain. Imaging Neuroscience, 2, 1–18. 10.1162/imag_a_00399

Gonzalez-Castillo, J., Saad, Z. S., Handwerker, D. A., Inati, S. J., Brenowitz, N., & Bandettini, P. A. (2012). Whole-brain, time-locked activation with simple tasks revealed using massive averaging and model-free analysis. Proceedings of the National Academy of Sciences of the United States of America, 109(14), 5487–5492. 10.1073/pnas.1121049109

Gutschalk, A., Hämäläinen, M. S., & Melcher, J. R. (2010). BOLD responses in human auditory cortex are more closely related to transient MEG responses than to sustained ones. Journal of Neurophysiology, 103(4), 2015–2026. 10.1152/jn.01005.2009

Handwerker, D. A., Ollinger, J. M., & D’Esposito, M. (2004). Variation of BOLD hemodynamic responses across subjects and brain regions and their effects on statistical analyses. NeuroImage, 21(4), 1639–1651. 10.1016/j.neuroimage.2003.11.029

Henson, R. N. A., Price, C. J., Rugg, M. D., Turner, R., & Friston, K. J. (2002). Detecting latency differences in event-related BOLD responses: Application to words versus nonwords and initial versus repeated face presentations. NeuroImage, 15(1), 83–97. 10.1006/nimg.2001.0940

Hu, S., Olulade, O., Castillo, J. G., Santos, J., Kim, S., Tamer, G. G., Luh, W. M., & Talavage, T. M. (2010). Modeling hemodynamic responses in auditory cortex at 1.5 T using variable duration imaging acoustic noise. NeuroImage, 49(4), 3027–3038. 10.1016/j.neuroimage.2009.11.051

Iglesias, J. E., Lorenzi, M., Ferraris, S., Peter, L., Modat, M., Stevens, A., Fischl, B., & Vercauteren, T. (2018). Model-Based Refinement of Nonlinear Registrations in 3D Histology Reconstruction. In A. F. Frangi, J. A. Schnabel, C. Davatzikos, C. Alberola-López, & G. Fichtinger (Eds.), Medical Image Computing and Computer Assisted Intervention – MICCAI 2018 (pp. 147–155). Springer International Publishing.

Josephs, O., Richardson, S., & Dick, F. (2025, September). High fidelity, reliable, and comfortable earbuds for auditory fMRI at all field strengths: OMEMS (Oto Micro Electrical Mechanical Systems). 8th International Conference on Auditory Cortex, Maastricht, NL.

Kenward, M. G., & Roger, J. H. (1997). Small sample inference for fixed effects from restricted maximum likelihood. Biometrics, 53(3), 983–997.

Kirilina, E., Lutti, A., Poser, B. A., Blankenburg, F., & Weiskopf, N. (2016). The quest for the best: The impact of different EPI sequences on the sensitivity of random effect fMRI group analyses. NeuroImage, 126, 49–59. 10.1016/j.neuroimage.2015.10.071

Kurzawski, J. W., Faruk Gulban, O., Jamison, K., & Winawer, J. (2022). Non-Neural Factors Influencing BOLD Response Magnitudes within Individual Subjects. Journal of Neuroscience, 42(38). 10.1101/2021.12.26.474185

Leung, K. K., Ridgway, G. R., Ourselin, S., & Fox, N. C. (2012). Consistent multi-time-point brain atrophy estimation from the boundary shift integral. NeuroImage, 59(4), 3995–4005. 10.1016/j.neuroimage.2011.10.068

Lewis, L. D., Setsompop, K., Rosen, B. R., & Polimeni, J. R. (2016). Fast fMRI can detect Oscillatory neural activity in humans. Proceedings of the National Academy of Sciences of the United States of America, 113(43), E6679–E6685. 10.1073/pnas.1608117113

Li, X., Morgan, P. S., Ashburner, J., Smith, J., & Rorden, C. (2016). The first step for neuroimaging data analysis: DICOM to NIfTI conversion. Journal of Neuroscience Methods, 264, 47–56. 10.1016/j.jneumeth.2016.03.001

Liégeois-Chauvel, C., Musolino, A., Badier, J. M., Marquis, P., & Chauvel, P. (1994). Evoked potentials recorded from the auditory cortex in man: evaluation and topography of the middle latency components. Electroencephalography and Clinical Neurophysiology/ Evoked Potentials, 92(3), 204–214. 10.1016/0168-5597(94)90064-7

Lindquist, M. A., Meng Loh, J., Atlas, L. Y., & Wager, T. D. (2009). Modeling the hemodynamic response function in fMRI: efficiency, bias and mis-modeling. NeuroImage, 45(1 Suppl). 10.1016/j.neuroimage.2008.10.065

López-Madrona, V. J., Trébuchon, A., Bénar, C. G., Schön, D., & Morillon, B. (2024). Different sustained and induced alpha oscillations emerge in the human auditory cortex during sound processing. Communications Biology, 7(1). 10.1038/s42003-024-07297-w

Lutti, A., Dick, F., Sereno, M. I., & Weiskopf, N. (2014). Using high-resolution quantitative mapping of R1 as an index of cortical myelination. In NeuroImage (Vol. 93, pp. 176–188). Academic Press Inc. 10.1016/j.neuroimage.2013.06.005

Mangal, J., Richardson, S., Brackenier, Y., Gardner, M., Di Cio, P., Casella, C., Malik, S., Hajnal, J., Callaghan, M. F., Dick, F., & Carmichael, D. W. (2025). Reducing motion artefact in high resolution 7T MRI using the Magnetic Resonance Minimal Motion (‘MR-MinMo’) head stabilisation device. 10.1101/2025.10.10.25337727

Moeller, S., Yacoub, E., Olman, C. A., Auerbach, E., Strupp, J., Harel, N., & Uǧurbil, K. (2010). Multiband multislice GE-EPI at 7 tesla, with 16-fold acceleration using partial parallel imaging with application to high spatial and temporal whole-brain FMRI. Magnetic Resonance in Medicine, 63(5), 1144–1153. 10.1002/mrm.22361

Moerel, M., De Martino, F., Kemper, V. G., Schmitter, S., Vu, A. T., Uğurbil, K., Formisano, E., & Yacoub, E. (2018). Sensitivity and specificity considerations for fMRI encoding, decoding, and mapping of auditory cortex at ultra-high field. NeuroImage, 164, 18–31. 10.1016/j.neuroimage.2017.03.063

Mottershead, J. P., Schmierer, K., Clemence, M., Thornton, J. S., Scaravilli, F., Barker, G. J., Tofts, P. S., Newcombe, J., Cuzner, M. L., Ordidge, R. J., McDonald, W. I., & Miller, D. H. (2003). High field MRI correlates of myelin content and axonal density in multiple sclerosis: A post-mortem study of the spinal cord. Journal of Neurology, 250(11), 1293–1301. 10.1007/s00415-003-0192-3

Patitucci, E. (2021). Investigating metabolic, vascular and structural neuroplasticity in healthy and diseased brain using advanced neuroimaging techniques.

Polimeni, J. R., & Lewis, L. D. (2021). Imaging faster neural dynamics with fast fMRI: A need for updated models of the hemodynamic response. In Progress in Neurobiology (Vol. 207). Elsevier Ltd. 10.1016/j.pneurobio.2021.102174

Posit team. (2025). RStudio: Integrated Development Environment for R.

Prince, J. S., Charest, I., Kurzawski, J. W., Pyles, J. A., Tarr, M. J., & Kay, K. N. (2022). Improving the accuracy of single-trial fMRI response estimates using GLMsingle. ELife, 11. 10.7554/eLife.77599

Proulx, S., Safi-Harb, M., LeVan, P., An, D., Watanabe, S., & Gotman, J. (2014). Increased sensitivity of fast BOLD fMRI with a subject-specific hemodynamic response function and application to epilepsy. NeuroImage, 93(P1), 59–73. 10.1016/j.neuroimage.2014.02.018

Robson, M. D., Dorosz, J. L., & Gore, J. C. (1998). Measurements of the Temporal fMRI Response of the Human Auditory Cortex to Trains of Tones.

Santoro, R., Moerel, M., De Martino, F., Goebel, R., Ugurbil, K., Yacoub, E., & Formisano, E. (2014). Encoding of Natural Sounds at Multiple Spectral and Temporal Resolutions in the Human Auditory Cortex. PLoS Computational Biology, 10(1). 10.1371/journal.pcbi.1003412

Schmierer, K., Wheeler-Kingshott, C. A. M., Tozer, D. J., Boulby, P. A., Parkes, H. G., Yousry, T. A., Scaravilli, F., Barker, G. J., Tofts, P. S., & Miller, D. H. (2008). Quantitative magnetic resonance of postmortem multiple sclerosis brain before and after fixation. Magnetic Resonance in Medicine, 59(2), 268–277. 10.1002/mrm.21487

Sereno, M. I., Sood, M. R., & Huang, R. S. (2022). Topological Maps and Brain Computations From Low to High. Frontiers in Systems Neuroscience, 16. 10.3389/fnsys.2022.787737

Siero, J. C. W., Petridou, N., Hoogduin, H., Luijten, P. R., & Ramsey, N. F. (2011). Cortical depth-dependent temporal dynamics of the BOLD response in the human brain. Journal of Cerebral Blood Flow and Metabolism, 31(10), 1999–2008. 10.1038/jcbfm.2011.57

Skipper, J. I., Van Wassenhove, V., Nusbaum, H. C., & Small, S. L. (2007). Hearing lips and seeing voices: How cortical areas supporting speech production mediate audiovisual speech perception. Cerebral Cortex, 17(10), 2387–2399. 10.1093/cercor/bhl147

Szycik, G. R., Stadler, J., Tempelmann, C., & Münte, T. F. (2012). Examining the McGurk illusion using high-field 7 tesla functional MRI. Frontiers in Human Neuroscience, (APRIL 2012). 10.3389/fnhum.2012.00095

Tabelow, K., Balteau, E., Ashburner, J., Callaghan, M. F., Draganski, B., Helms, G., Kherif, F., Leutritz, T., Lutti, A., Phillips, C., Reimer, E., Ruthotto, L., Seif, M., Weiskopf, N., Ziegler, G., & Mohammadi, S. (2019). hMRI – A toolbox for quantitative MRI in neuroscience and clinical research. NeuroImage, 194, 191–210. 10.1016/j.neuroimage.2019.01.029

Taylor, A. J., Kim, J. H., & Ress, D. (2018). Characterization of the hemodynamic response function across the majority of human cerebral cortex. NeuroImage, 173, 322–331. 10.1016/j.neuroimage.2018.02.061

Taylor, A. J., Kim, J. H., & Ress, D. (2022). Temporal stability of the hemodynamic response function across the majority of human cerebral cortex. Human Brain Mapping, 43(16), 4924–4942. 10.1002/hbm.26047

Triantafyllou, C., Hoge, R. D., Krueger, G., Wiggins, C. J., Potthast, A., Wiggins, G. C., & Wald, L. L. (2005a). Comparison of physiological noise at 1.5 T, 3 T and 7 T and optimization of fMRI acquisition parameters. NeuroImage, 26(1), 243–250. 10.1016/j.neuroimage.2005.01.007

Triantafyllou, C., Hoge, R. D., Krueger, G., Wiggins, C. J., Potthast, A., Wiggins, G. C., & Wald, L. L. (2005b). Comparison of physiological noise at 1.5 T, 3 T and 7 T and optimization of fMRI acquisition parameters. NeuroImage, 26(1), 243–250. 10.1016/j.neuroimage.2005.01.007

Watanabe, M., Bartels, A., Macke, J., Murayama, Y., & Logothetis, N. (2013). Temporal Jitter of the BOLD Signal Reveals a Reliable Initial Dip and Improved Spatial Resolution. Current Biology, 23(21), 2146–2150. 10.1016/j.cub.2013.08.057

Weiskopf, N., Suckling, J., Williams, G., Correia M., M. M., Inkster, B., Tait, R., Ooi, C., Bullmore T., E. T., & Lutti, A. (2013). Quantitative multi-parameter mapping of R1, PD*, MT, and R2* at 3T: A multi-center validation. Frontiers in Neuroscience, (7 JUN). 10.3389/fnins.2013.00095

Worsley, K. J., Liao, C. H., Aston, J., Petre, V., Duncan, G. H., Morales, F., Evans, A. C., & Worsley, K. (2002). A General Statistical Analysis for fMRI Data.

Yeşilyurt, B., Uǧurbil, K., & Uludaǧ, K. (2008). Dynamics and nonlinearities of the BOLD response at very short stimulus durations. Magnetic Resonance Imaging, 26(7), 853–862. 10.1016/j.mri.2008.01.008

Zaidi, A. D., Birbaumer, N., Fetz, E., Logothetis, N., & Sitaram, R. (2023). The hemodynamic initial-dip consists of both volumetric and oxymetric changes reflecting localized spiking activity. Frontiers in Neuroscience, 17. 10.3389/fnins.2023.1170401

Zeidman, P., Kazan, S. M., Todd, N., Weiskopf, N., Friston, K. J., & Callaghan, M. F. (2019). Optimizing data for modeling neuronal responses. Frontiers in Neuroscience, 13(JAN). 10.3389/fnins.2018.00986

Zhang, N., Yacoub, E., Zhu, X. H., Ugurbil, K., & Chen, W. (2009). Linearity of blood-oxygenation-level dependent signal at microvasculature. NeuroImage, 48(2), 313–318. 10.1016/j.neuroimage.2009.06.071

Zulfiqar, I., Havlicek, M., Moerel, M., & Formisano, E. (2021). Predicting neuronal response properties from hemodynamic responses in the auditory cortex. NeuroImage, 244. 10.1016/j.neuroimage.2021.118575

